# AI recognizes convergent somatic hypermutation signatures to allow the discovery of variant-resilient broadly neutralizing antibodies

**DOI:** 10.64898/2025.12.10.693372

**Authors:** Qihong Yan, Xiaohan Huang, Banghui Liu, Yicheng Yao, Huiran Zheng, Yidong Song, Fandi Wu, Zhixiao He, Shaowei Li, Fangli Chen, Chuanying Niu, Zimu Li, Hang Yuan, Yuting Lin, Lan Chen, Na Zang, Juxue Xiao, Tingting Wang, Canjie Chen, Haoshi Bai, Jia Li, Mengyu Wu, Jingxian Zhao, Jianhua Yao, Yuedong Yang, Airu Zhu, Xiaoli Xiong, Jincun Zhao

## Abstract

Elucidating the mechanisms by which broadly neutralizing antibodies (bnAbs) develop to confer durable immunity in humans is pivotal for the rational design of next-generation vaccines and therapeutics. VH3-53/3-66-encoded public antibodies are widely elicited in the population after the COVID-19 pandemic. Here, we isolated 15 VH3-53/3-66-encoded bnAbs from elite neutralizers who experienced sequential SARS-CoV-2 Omicron breakthrough infections. These bnAbs exhibit exceptional potency and breadth against circulating variants and, as members of a long-lived public antibody lineage, likely contribute to durable humoral protection. Genetic analysis of VH3-53/3-66-encoded bnAbs reveals a convergent somatic hypermutation pattern at seven positions in and around the CDR loops that emerges with repeated viral exposures and distinguishes them from non-bnAbs. Grafting these combined mutations onto clinically escaped antibodies broadens their breadth and restores neutralization activity. Structural analysis shows that these convergent mutations cooperatively remodel CDRs to bind and tolerate virus receptor-binding domain mutations. Using an AI-based antibody language model trained on data linking somatic hypermutations to binding affinity and neutralization, the model is able to identify mutational patterns predictive of breadth, enabling the discovery of a rare bnAb with protective activity against the latest variants from early-pandemic antibody repertoires. This demonstrates the feasibility of establishing an AI-empowered pipeline to identify mutation-tolerant bnAbs from early-pandemic antibody repertoires to fight fast-evolving newly introduced viruses.

## Introduction

Broadly neutralizing antibodies (bnAbs) have shown promises in combating highly mutable RNA viruses such as influenza, HIV, and SARS-CoV-2^1–3^. The capacity of broadly neutralizing antibodies (bnAbs) to neutralize diverse viral strains makes them invaluable guides for vaccine and therapeutic design; however, their rational elicitation or discovery remains a major challenge. For most RNA viruses, bnAbs emerge only after multiple antigen exposures and prolonged affinity maturation, and they typically constitute an exceedingly small fraction of the overall antibody repertoire^4^. Moreover, the molecular rules that govern their development remain incompletely understood, severely constraining efforts to generate bnAbs early during outbreaks of emerging pathogens.

During the COVID-19 pandemic, the continued and rapid emergence of immune escape variants has highlighted these limitations. Despite intensive global vaccination and infection-driven immunity, the establishment of long-lasting, cross-variant neutralizing antibody responses has remained challenging. Shared (“public”) neutralizing antibodies targeted immunodominant epitopes on the spike receptor-binding domain (RBD) have been widely identified from convalescents or vaccinee donors, particularly those encoded by VH1-2, VH1-58, VH1-69, and VH3-53/3-66 germlines^5–7^. Among these, VH3-53/3-66 antibodies are the most prevalent and target a class 1 RBD epitope with potent neutralization against the ancestral strain^8^. However, this immunodominant class has also been highly vulnerable to antigenic drift. VH3-53/3-66 antibodies isolated early in the pandemic, such as B-38^9^, COVA2-04^10^, CC12.1^11^, and clinically approved LY-CoV016 (CB6)^12^, P2C-1F11^13^, and BD-604^14^, were rapidly escaped by variants carrying K417N or K417T substitutions that are features of Beta, Gamma, and Omicron sublineages^11,15,16^. Intriguingly, despite extensive viral evolution, a few isolated VH3-53/3-66 antibodies retain cross-variant neutralizing activity. They appear to achieve this through the acquisition of somatic hypermutations (SHMs) that optimize binding to divergent RBDs^17–19^. Longitudinal analyses of convalescents and vaccinees reveal that persistent antigenic stimulation, through prolonged viral RNA or antigen presence, as well as sequential variant exposures^20^, drives continued affinity maturation of pre-existing VH3-53/3-66 memory B cells^21^. This process promotes somatic hypermutations associated with expanded antibody breadth and increased resilience to viral escape^22,23^. Some evidence indicates that matured VH3-53/3-66 antibodies carry stereotypic SHMs such as F27V/I, T28I, S31R, and Y58F, which remodel the paratope to tolerate RBD variation^19,24,25,26–28^. BD55-1205, for instance, a VH3-66-encoded neutralizing antibody isolated from a WT SARS-CoV-2 convalescent, maintains potent neutralization against all major circulating variants through three unique residues (R31, P53, and R102) in its heavy-chain complementarity-determining regions (CDRs) that introduce additional polar interactions at the binding interface^19^. It remains unclear whether convergent somatic hypermutation (SHM) trajectories drive the natural evolution of broad-spectrum VH3-53/3-66 antibodies. Defining such convergent pathways may reveal how cross-variant breadth is acquired and provide a molecular blueprint for broadly neutralizing antibody generation.

The ability to rapidly identify bnAbs that can withstand future antigenic drift represent a major unmet need in pandemic preparedness. The pace of therapeutic antibody discovery still lags behind viral evolution, despite remarkable progress in antibody engineering. The rapid advancement of artificial intelligence (AI), particularly protein language models (PLMs) trained on large-scale immunoglobulin repertoires, offers a potential transformative route to close this gap^24,29–31^. These models can learn the intrinsic grammar of antibody sequence evolution, enabling fitness prediction, structure inference, and sequence generation with unprecedented speed and precision^32–35^.

Here, we mechanistically characterize fifteen VH3-53/3-66-encoded, broad-spectrum human monoclonal antibodies isolated from elite neutralizers with repeated Omicron breakthrough infections. These antibodies exhibit exceptional potency and breadth across all major SARS-CoV-2 variants. Integrated genetic and structural analyses show that repeated Omicron exposures drive convergent somatic hypermutations (SHMs) that expand neutralization breadth; introducing these SHMs into clinically escaped VH3-53/3-66 antibodies restores neutralization.

We reasoned that the convergent SHM patterns observed in naturally matured VH3-53/3-66 antibodies encapsulate the molecular logic of bnAb derivation. Accordingly, we show that an antibody language model trained on SHM-informed repertoires can learn mutational signatures predictive of breadth, enabling in silico identification of potent bnAbs before their mutation-tolerant features are experimentally selected. This strategy yields the AI-identified bnAb R102-9, which neutralizes all circulating variants and confers robust protection in vivo. By coupling SHM-convergence analysis with AI-driven inference, we establish a predictive antibody discovery workflow that transforms bnAb discovery from retrospective observation into a proactive, data-guided process.

## Results

### SARS-CoV-2 spike accumulates sequential mutations on the VH3-53/3-66-targeted epitope

Canonical class 1 VH3-53/3-66 antibodies neutralize SARS-CoV-2 by blocking ACE2-RBD binding (Figure 1A). Structural analysis identified 20 amino acid residues on the WT RBD that directly participate in ACE2 interaction (Figure 1B). To investigate immune evasion of VH3-53/3-66 antibodies by viral variants, we analyzed epitope profiles of 79 structurally characterized VH3-53/3-66 antibodies, revealing remarkable epitope convergence (Figure S1). Our comprehensive mapping identified 28 RBD residues exhibiting substantial antibody interactions (defined by mean buried surface area >10 Å^2^; Figure 1C). Evolutionary tracking across major circulating variants identified 34 mutated sites within the RBD (Figure 1D). Notably, we observed 10 overlapping residues among: (1) ACE2-binding sites, (2) VH3-53/3-66 epitopes, and (3) variant mutation hotspots (Figure 1E). Nine of the ten overlapping residues (R403, K417, L455, F456, N460, A475, F486, Q493, and N501) have been experimentally validated to mediate different levels of neutralization escape^25,36–38^. Strikingly, phylogenetic mapping revealed progressive accumulation of these escape mutations across successive variants within the VH3-53/3-66 epitope (Figure 1F). The earliest emerged B.1.1.7 (Alpha) variant carried only the N501Y mutation, followed by B.1.351 (Beta) acquiring K417N at the epitope core. Omicron BA.1/BA.2 acquired Q493R over Beta’s profile while BA.4/BA.5 gained F486V. BQ.1 and XBB evolved N460K on BA.4/5 backgrounds, with XBB initially carrying F486S before its descendants acquired F486P. Subsequent XBB-derived lineages (EG.5, HV.1, and HK.3) sequentially acquired F456L and L455F. The XBB sub-lineage JD.1.1 emerged with the most extensive set of seven mutations including distinctive A475V. BA.2.86 emerged with five mutations within the VH3-53/3-66-targeted epitope, followed by JN.1 and XDV.1 evolved L455S and F456L; while the recently circulating KP.3.1.1, LP.8.1 and the currently dominant NB.1.8.1 all acquired Q493E, culminating in an eight-mutation escape profile (Figure 1F). Early in the pandemic, the K417N mutation was the primary signature identified to confer the escape of VH3-53/3-66 antibodies; however, our analysis demonstrates a continued accumulation of escape mutations within the VH3-53/3-66 epitope. The shift from isolated single-amino acid changes early in the pandemic to multi-mutation constellations underscores sustained evolutionary pressure on this epitope. Therefore, we hypothesize that pre-existing memory B cells utilizing VH3-53/3-66 are still capable of cross-recognizing their convergent epitope even in the most recent SARS-CoV-2 variants.

**Figure 1.**
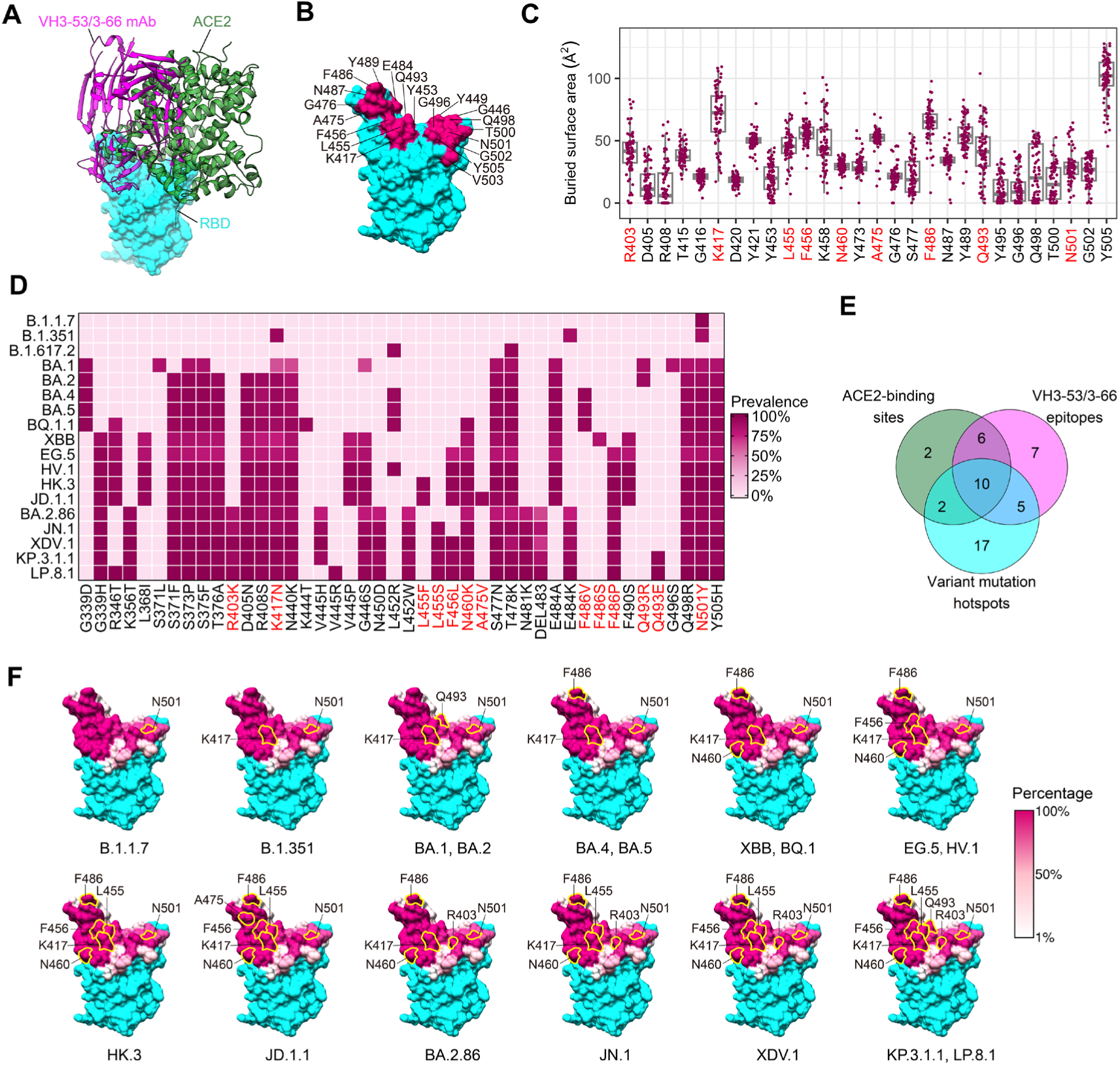
SARS-CoV-2 spike accumulates sequential mutations on the VH3-53/3-66-targeted epitope. (**A**) Comparison of VH3-53/3-66 mAbs (BD-604 as representative, PDB: 7CHF) and ACE2 (PDB: 6M0J) binding to the RBD. VH3-53/3-66 mAbs, ACE2 and RBD are colored in magenta, green and cyan, respectively. (**B**) The epitope residues of ACE2 binding sites, binding interface of ACE2 highlighted in magenta. (**C**) The buried surface area of 79 VH3-53/3-66 mAbs at each binding sites. Nine residues that have been experimentally validated to mediate antibody neutralization escape were highlighted with red. (**D**) The mutation profile on RBD of 18 SARS-CoV-2 variants. Nine residues that have been experimentally validated to mediate antibody neutralization escape were highlighted with red. (**E**) Venn-plot showing the overlapped residues among ACE2-binding sites, VH3-53/3-66 epitopes, and variant mutation hotspots. (**F**) Accumulation of mutations within the epitope of VH3-53/3-66 mAbs across SARS-CoV-2 variants. Footprints of VH3-53/3-66 epitopes on RBD, and the key escape mutations within the epitope was labeled. Epitope residues on the RBD are colored according to the proportion of VH3-53/3-66 mAbs interacting with each positions.

### The VH3-53/3-66 antibody lineage constitutes a predominant component of durable humoral immunity in elite neutralizers

To efficiently isolate VH3-53/3-66 antibodies with broad neutralization breadth and tolerance to variant escape, we focused on screening elite neutralizers for monoclonal antibody isolation. From a cohort of 68 convalescent individuals with three doses of inactivated vaccine followed by BA.5 and XBB breakthrough infections, we identified 4 subjects with high neutralizing titers (FRNT_50_ > 2,000) against WT, BA.5, XBB.1, and EG.5 strains (Figure S2A). Using Avi-tag-biotinylated JD.1.1 and KP.3.1.1 RBD probes, we isolated a total of 3715 JD.1.1/KP.3.1.1-specific memory B cells through fluorescence-activated cell sorting (FACS) (Figure S2B). We profiled sorted cells by 5’ directed scRNA-seq for both paired V(D)J and mRNA sequencing, recovering matched single-cell V(D)J and transcriptome profiles in 2124 cells. According to their transcriptomes, cells were clustered into four distinct clusters by Uniform Manifold Approximation and Projection for Dimension Reduction (UMAP) (Figure 2A-B). We also quantified their antibody isotype frequency and SHM levels (Figure 2C-D). Cluster 1 expressed Ig genes with little to no SHM or CSR (class switch recombination) and gene signatures associated with naive B cells (*TCL1A, BACH2, IGHD*), suggesting that this subset was composed of naive-like B cells or very recently activated B cells (*CD69, CD83*) (Figure 2B-D). Clusters 2-4 with patterns of higher SHM and CSR were charactered as memory B cells (MBCs) (Figure 2B-D). The classical MBC gene signature (*CD27*) was mapped to clusters 2 and 3 (Figure 2B). However, cluster 2 contained higher frequency of un-class-switched B cells with lower SHM levels, while cluster 3 were highly mutated and were largely class-switched to IgG1 subclass (Figure 2C-D). Cluster 4 exhibited high expression of *ITGAX* (encoding CD11c) and *ZEB2*, both of which are strongly linked to double-negative 2 (DN2) B cells, as well as the inhibitory receptor gene *FCRL5* (Fc receptor-like 5) (Figure 2B).

**Figure 2.**
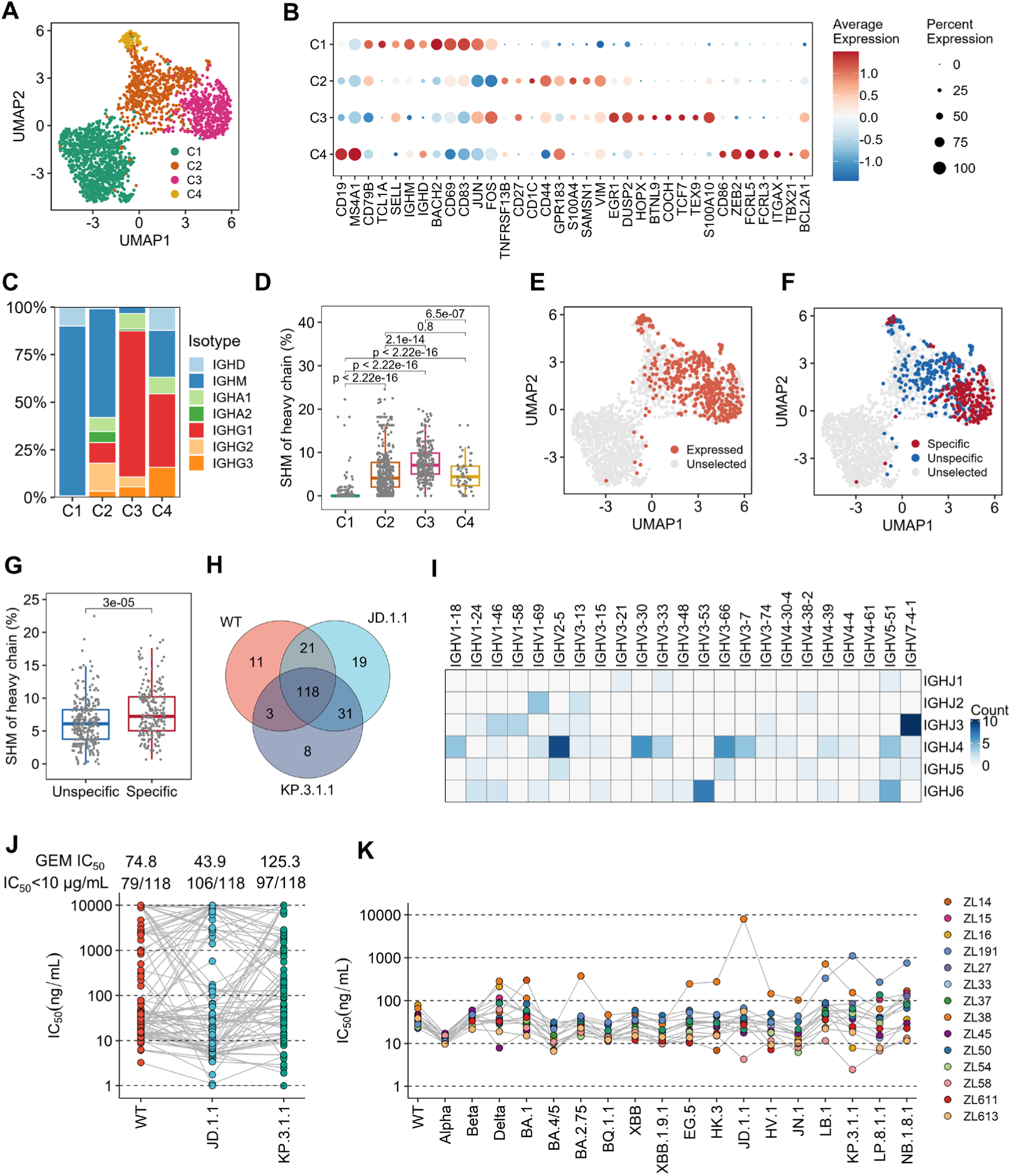
Isolation of broad neutralizing mAbs from SARS-CoV-2 elite neutralizers. (**A**) Uniform manifold approximation projection (UMAP) embedding of scRNA-seq transcriptome profiles of the 2124 JD.1.1 and KP.3.1.1 probe+ B cells, colored by cluster. (**B**) Dotplot showing the marker genes of the four B cell clusters. (**C**) Comparison of isotype frequency of the four B cell clusters. (**D**) Comparison of somatic hypermutation rates among the four B cell clusters. (**E**) UMAP highlighting cells from which mAb was synthesized and expressed. The synthesized and expressed mAbs are colored with red dots and unselected mAbs with grey dots. (**F**) UMAP showing the RBD-specific mAbs with red dots, RBD-unspecific mAbs with blue dots, and unselected mAbs with grey dots. (**G**) Comparison of somatic hypermutation rates between RBD-specific and unspecific mAbs. (**H**) Venn-plot showing the number of mAbs with WT, JD.1.1 and KP.3.1.1 cross-reactivity. (**I**) IGHV and IGHJ germline gene usage of the 118 mAbs with WT, JD.1.1 and KP.3.1.1 cross-reactivity. (**J**) Neutralization activities of the 118 mAbs to WT, JD.1.1 and KP.3.1.1. Geometric mean of IC_50_ values and numbers of nAbs (IC_50_ < 10 μg/mL) are annotated above each group of points. (**K**) Neutralization activities of the 15 VH3-53/3-66 mAbs to SARS-CoV-2 variants.

We synthesized and expressed 486 mAbs from none-IGHD and none-IGHM-expressing B cells to increase the chance of identifying neutralizing antibodies. These B cells represent nearly 23% of all B cells obtained by scRNA-seq (Figure 2E). Totally, 207 of the selected mAbs from cells isolated using JD.1.1/KP.3.1.1 probes exhibited specific binding to at least one tested RBD (WT, JD.1.1, or KP.3.1.1) by biolayer interferometry (BLI) (Figure 2F). Notably, more than 91% (189/207) of the RBD-specific B cells are identified within the classical MBC cluster 3 (Figure 2F), which displays the highest level of somatic hypermutation (SHM) among all subsets (Figure 2D). Moreover, *BCL2A1*, a key anti-apoptotic regulator, is highly expressed in cluster 3, suggesting this B cell subset may underlie long-lived humoral immunity (Figure 2B). The SHM levels were significantly higher in RBD-specific monoclonal antibodies compared to unspecific counterparts (Figure 2G). Out of the 207 RBD-specific antibodies, a total of 118 candidates achieved binding in BLI assays against WT, JD.1.1, and KP.3.1.1 RBD at the same time (Figure 2H). Analysis of germline genes showed that *IGHV2-5*, *IGHV3-30*, *IGHV3-53*, *IGHV3-66*, *IGHV5-51*, and *IGHV7-4-1* for the heavy chain were predominantly used among the 118 cross-reactive mAbs (Figure 2I). To narrow the range of candidates, we evaluated their neutralizing activities against WT, JD.1.1, and KP.3.1.1 SARS-CoV-2 pseudoviruses. Although these mAbs were isolated using JD.1.1/KP.3.1.1 probes, 67% (79/118) of which showed effective neutralization with a geometric mean 50% inhibitory concentration (GEM IC_50_) of 74.8 ng/mL against WT (Figure 2J), indicating that most of these were derived from recalled memory B cells. Finally, we identified 60 mAbs that could effectively neutralize three strains (Figures 2J, S2C, Table S1). We next tested if the cross-neutralizing mAbs were enriched for Ig genes VH3-53 and VH3-66, features that are preferentially found among class 1 nAbs. Our analysis revealed that 15 (25%) of the 60 cross-neutralizing antibodies utilized the VH3-53/3-66 gene segments, representing the second most prevalent genotype after *IGHV7-4-1* (Figure S2C). In addition, all 15 VH3-53/3-66-derived antibodies exhibited canonical class 1 characteristics, featuring short HCDR3 loops of 8-12 amino acids in length (Figure S2D). To investigate the breadth of 15 VH3-53/3-66-derived antibodies, we tested their neutralization activities against 19 representative SARS-CoV-2 pseudoviruses. Notably, 13 of the 15 VH3-53/3-66 antibodies demonstrated potent neutralization against all 19 pseudoviruses tested, with IC_50_ values <50 ng/mL for most strains (Figure 2K). The exceptions were ZL38 and ZL191, which showed reduced efficacy against the emerging JD.1.1, LB.1, KP.3.1.1, LP.8.1.1 and NB.1.8.1 variants (Figure 2K). Interestingly, four of the 15 VH3-53/3-66-class mAbs were derived from IGHA1^+^ B cells, suggesting preferential induction of these broadly neutralizing antibodies at mucosal sites (Figure S2D). Taken together, we identified 15 V3-53/3-66 public antibodies with broad neutralizing capacity against SARS-CoV-2, which dominate the long-lived memory B cell population and exhibit a mucosa-associated antibody isotype expression, suggesting their dual origin from systemic and mucosal immune responses.

### Repeated Omicron breakthrough infections drive convergent somatic hypermutations within VH3-53/3-66 antibodies

We next directly compared the *in vitro* neutralization potency and breadth of the 15 mAbs with the previously published VH3-53/3-66 antibodies, using a diverse panel of strains from XBB and JN.1 sub-lineages (Figure 3A). A panel of 32 representative VH3-53/3-66 neutralizing antibodies derived from individuals with different exposure histories were included. Overall, antibodies derived from either wild-type (WT) infection or vaccine recipients exhibited the most restricted neutralization breadth, with a median 50% inhibitory concentration (median IC_50_) of 61.78-10000 ng/mL. In contrast, monoclonal antibodies that isolated from individuals with single Omicron breakthrough infection demonstrated improved cross-neutralization, with median IC_50_ of 33.21-1315 ng/mL. Most notably, antibodies that we isolated from elite neutralizers with sequential BA.5 and XBB breakthrough infections displayed the most exceptional pan-variant neutralization breadth, with median IC_50_ of 14.57-112.8 ng/mL (Figure 3A, Table S2). Among all the previously published antibodies tested, BD55-1205 is the only one that demonstrated neutralizing activity against all the viruses tested with a median IC_50_ of 485.1 ng/ml (range 52.35-791.7 ng/mL) (Figure 3A). These findings suggest that repeated heterologous antigen exposure significantly promotes the development of SARS-CoV-2 broadly neutralizing antibodies, a process we propose to be closely associated with the accumulation of somatic hypermutations.

**Figure 3.**
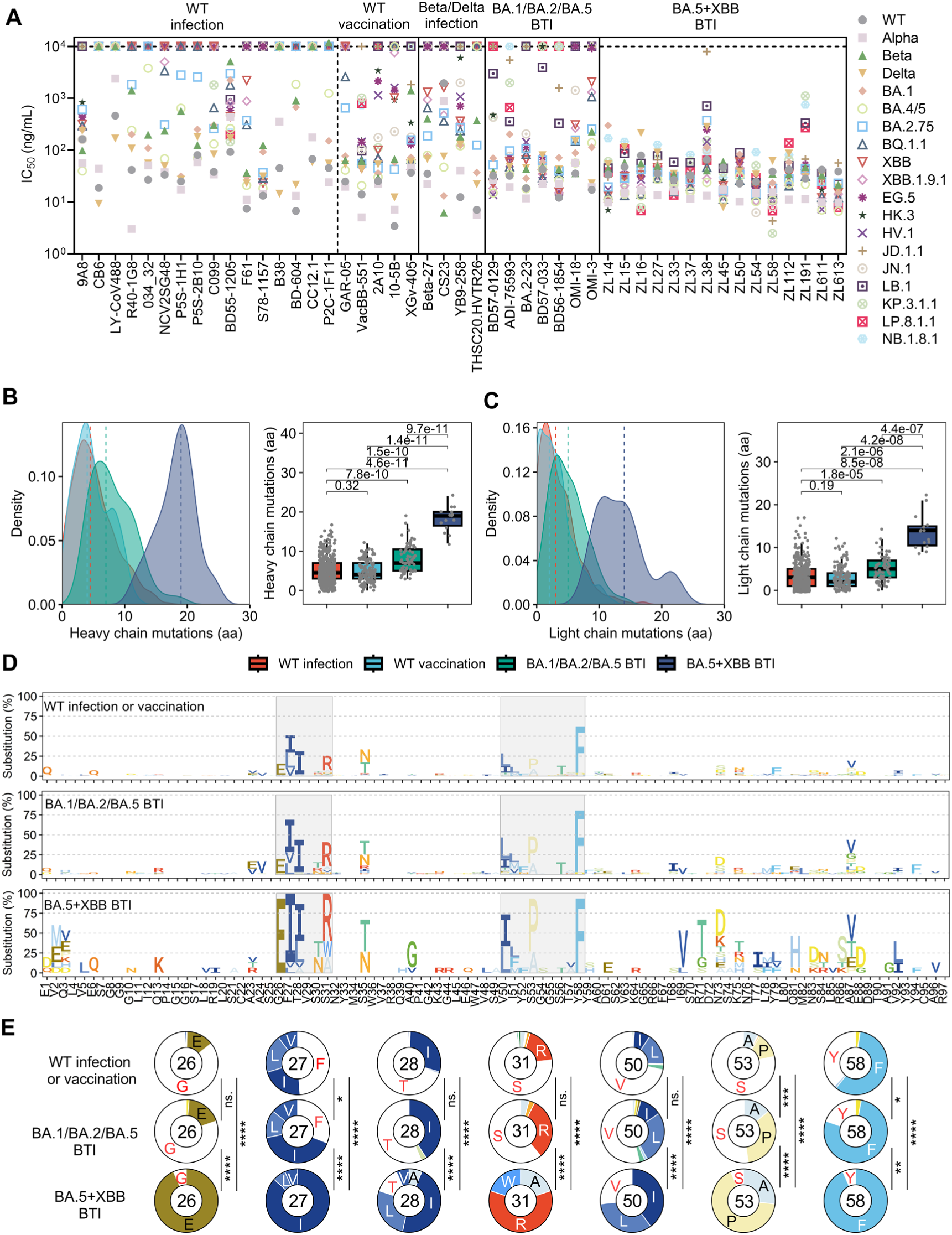
Comparison of the neutralizing activities and somatic hypermutations of VH3-53/3-66 mAbs. (**A**) Comparison of neutralizing activities (IC_50_) of VH3-53/3-66 mAbs isolated from different infection or vaccination background. BTI, breakthrough infection. (**B-C**) Comparison of somatic hypermutation levels within heavy (**B**) or light (**C**) chain variable region of VH3-53/3-66 mAbs isolated from different infection or vaccination background. (**D**) Somatic mutation profiles of VH3-53/3-66 mAbs isolated from different infection or vaccination background. (**E**) Pie charts indicate the frequency of germline (white) and mutated residues at position 26, 27, 28, 31, 50, 53, and 58 of heavy chain variable region of VH3-53/3-66 mAbs.

To understand the development of broad neutralizing activity, we further characterized the somatic hypermutation features of these 15 mAbs compared to other published VH3-53/3-66 public antibodies. We collected a total of 708 published VH3-53/3-66 antibodies that characterized as class 1 features from CoV-AbDab^39^. As previously reported, VH3-53/3-66 antibodies isolated early in the pandemic generally have no to limited mutations, with more than 60% antibodies have 0–5 mutations and less than 10% antibodies have more than 10 mutations (Figures 3B-C, S3A). Following single Omicron BA.1/BA.2/BA.5 breakthrough infection (BTI), VH3-53/3-66 antibodies exhibited significantly increased SHMs in both heavy and light chains. Repeated Omicron breakthrough infection (BA.5+XBB BTI) induced substantial further accumulation of SHMs (Figures 3B-C, S3A).

To investigate potential enrichment of specific SHMs, we analyzed the mutation pattern at each residue position across the FR1-FR3 regions of VH3-53/3-66 antibody heavy chains. Notably, we identified seven prominent convergent somatic mutations in antibody CDR1 and CDR2 regions, with this enrichment pattern becoming more pronounced following both primary and secondary Omicron breakthrough infections (Figure 3D-E). The convergent SHM profile comprised both some of previous documented substitutions F27I, T28I, S31R, V50L, S53P, and Y58F, as well as a newly observed prevalent mutation G26E. Notably, although convergent somatic hypermutation patterns become apparent when antibody sequences induced by WT infection or vaccination are cross-compared, these mutations rarely co-occur within the same antibody that was isolated early in the pandemic (Figures 3D, S3B). Following two Omicron breakthrough infections, we observed a marked increase in the proportion of VH3-53/3-66 antibodies harboring 6-7 concurrent somatic mutations (Figure 3E, S3B). Strikingly, all 15 neutralizing antibodies isolated in this study exhibited near-complete convergence at the seven critical paratope residues, indicative of strong antigen-driven selection during affinity maturation (Figure S3B). Furthermore, we analyzed the paired light chain repertoire of VH3-53/3-66 antibodies, revealing an enrichment of VK1-33 following breakthrough infections (Figure S3C). Using the same analytical approach, we identified three convergent SHMs (30_L_-32_L_) clustered together within the LCDR1 paratope of VK1-33-encoded light chains (P<0.001 for each site, χ² test) (Figure S3D-E). Taken together, these results demonstrate that repeated antigen exposure markedly enriches convergent somatic mutations in both heavy and light chains of VH3-53/3-66 mAbs.

### Evolution selected convergent somatic hypermutations expand potency and breadth of VH3-53/3-66 antibodies

To precisely define the function of the convergent SHMs in the isolated cross-neutralization of antibodies, we reverted the heavy chains of the matured ZL54, ZL58, ZL112, and ZL611 antibodies to their germline forms (ZL54gl, ZL58gl, ZL112gl, and ZL611gl). While the germline versions could neutralize WT, albeit 30.3- to 83.5-folds less potently than the mature antibodies, they lacked neutralizing activity against the tested variants (Figures 4A, S4A), confirming the essential role of affinity maturation introduced residues for neutralization breadth. To understand the roles of the newly observed novel dominant mutation G26E, we reverted 26E to a germline 26G in matured ZL54, ZL58, ZL112, and ZL611 (Figure 4A-B). Neutralization assays demonstrated that four engineered antibody mutants (ZL54-E26G, ZL58-E26G, ZL112-E26G, and ZL611-E26G) showed compromised neutralization potency against most circulating SARS-CoV-2 variants compared to their parental antibodies (2.5- to 10.3-fold increase in mean IC_50_; P < 0.01). Most notably, neutralization was most severely impaired against recently emerging Omicron subvariants, including KP.3.1.1, LP.8.1.1 and NB.1.8.1 (3.5- to 72-fold IC_50_ increase), highlighting G26E’s role in conferring continued cross-neutralization (Figures 4A-B, S4B-D).

**Figure 4.**
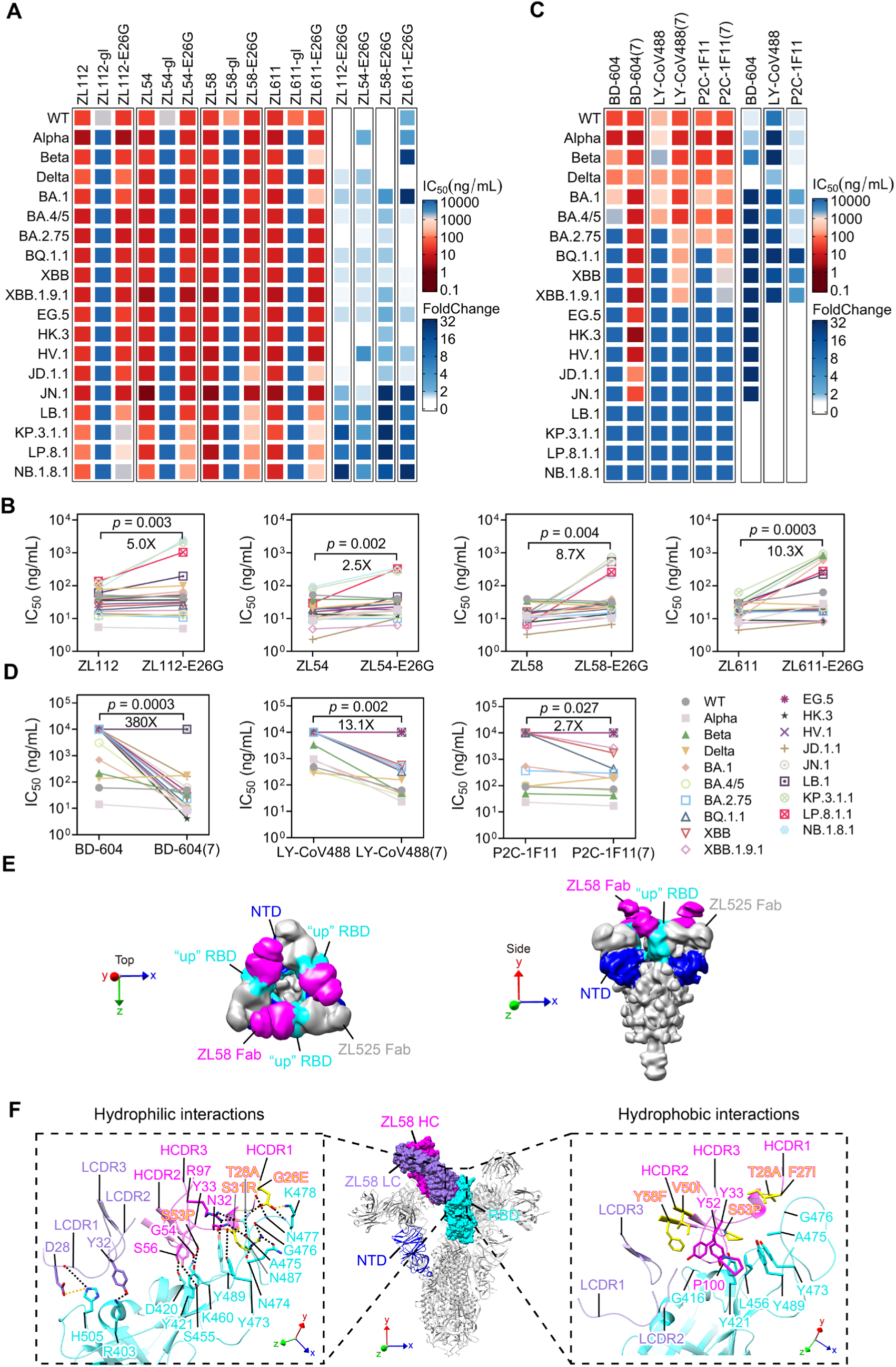
Convergent SHM extent the neutralization and potency and breadth of VH3-53/3-66 mAbs. (**A**) Heatmap (left panel) showing the neutralizing activity of ZL112, ZL54, ZL58, and ZL611, as well as their mutants against 19 major SARS-CoV-2 strains. Heatmap (right panel) showing the foldchange of IC_50_ of E26G versions compared to ZL112, ZL54, ZL58, and ZL611, respectively. (**B**) Comparison of the IC_50_ values between ZL112, ZL54, ZL58, or ZL611 and their E26G mutants. (**C**) Heatmap (left panel) showing the neutralizing activity of BD-604, LY-CoV488, and P2C-1F11, as well as their mutants against 19 major SARS-CoV-2 strains. Heatmap (right panel) showing the foldchange of IC_50_ of SHM-introduction versions compared to BD-604, LY-CoV488, and P2C-1F11, respectively. (**D**) Comparison of the IC_50_ values between BD-604, LY-CoV488, and P2C-1F11 and their SHM-introduction mutants. (**E**) Structures (low-pass filtered to 12 Å) of Omicron LP.8.1 spike (S-6P) in complex with ZL58 Fab and ZL525 Fab. ZL58 Fab, NTD and RBD are highlighted in magenta, blue and cyan, respectively; the rest of the spike and ZL525 Fab are colored gray. (**F**) Epitope of ZL58 on the RBD of Omicron LP.8.1 spike. Detailed interactions between ZL58 and RBD are shown in the dashed boxes. CDR loops are indicated, and selected interacting residues in antibody-RBD interface are shown. Somatically hypermutated residues are highlighted in yellow and labeled with the changes indicated. Backbone carbonyl oxygens and amide nitrogens are indicated by red and blue dots, respectively. Yellow and black dashed lines indicate salt bridge and hydrogen bond interactions, respectively.

Oppositely, we introduced the seven somatic hypermutations into the germline version of previously clinically approved mAbs BD-604, P2F-1C11, and LY-CoV488, to generate their artificially affinity matured versions designated as BD-604(7), P2F-1C11(7), and LY-CoV488(7), to investigated whether the identified SHM signature would confer increased breadth to neutralize Omicron subvariants (Figure 4C). Neutralization results show that these specific mutations indeed conferred the three antibodies with enhanced neutralizing activity against Omicron subvariants, including BA.1, BA.4/5, BA.2.75, BQ.1.1, XBB.1.9.1 (Figure 4C). Of note, these mutations even enabled BD-604 to neutralize EG.5, HK.3, HV.1, JD.1.1 and JN.1. Overall, the neutralizing activity of the artificially affinity matured mAbs display 380-, 13.1-, 2.7-folds improvement compared to their parent antibodies, respectively (Figure 4D). These results show that the efficacy of clinically escaped therapeutics antibodies can be restored by the introduction of the seven convergent hotspot mutations, increasing their neutralizing breadth.

To structurally define the contribution of SHM-introduced residues to RBD binding by VH3-53/3-66 antibodies, we determined a cryo-EM structure of the Omicron LP.8.1.1 spike (S-6P) in complex with the ZL58 Fab, a representative VH3-53/3-66 antibody. We included the ZL525 Fab, an R1-32-like public antibody known to induce RBD opening^7^, to enhance RBD structural resolution through increased molecular mass. We only observed the 1:3:3 S-trimer:ZL58:ZL525 complex, with all three RBDs adopting an “up” conformation (Figures 4E, S5-6). Based on the locally refined RBD-Fab map at 2.93 Å resolution (Figures S5-6), we built an atomic model of the complex, enabling detailed characterization of the ZL58 binding interface (Figure 4F). As expected, ZL58 is a classical class 1 antibody that binds to the apical head of RBD, overlapping the receptor-binding motif. Buried surface area analysis reveals that ZL58 engages the RBD primarily through HCDR1, HCDR2, HCDR3, LCDR1, and LCDR3, with the heavy and light chains contributing 689 Å² and 192 Å², respectively. The binding interface consists of a network of hydrophilic and hydrophobic interactions. Hydrophilic contacts, including 16 hydrogen bonds and 2 salt bridges, involve RBD residues R403_S_, D420_S_, Y421_S_, S455_S_, K460_S_, Y473_S_, N474_S_, A475_S_, G476_S_, N477_S_, K478_S_, N487_S_, Y489_S_, and H505_S_ that interact with HCDR residues E26^H^, A28^H^, R31^H^, N32^H^, Y33^H^, P53^H^, G54^H^, S56^H^, R97^H^, and LCDR residues D28^L^ and Y32^L^. Hydrophobic contacts involve RBD residues G416_S_, Y421_S_, L456_S_, Y473_S_, A475_S_, G476_S_, and Y489_S_ packing against HCDR residues I27^H^, A28^H^, Y33^H^, Y52^H^, P53^H^, F58^H^, and P100^H^. Notably, many of these interactions are mediated by residues introduced through somatic hypermutation, including E26^H^, I27^H^, A28^H^ and R31^H^ in HCDR1, and P53^H^ and F58^H^ in HCDR2, underscoring the indispensable role of SHM in developing ZL58’s breadth. In particular, the newly observed dominant mutation G26E^H^ introduced a salt bridge to K478_S_, and a network of charged hydrogen bonding interactions at the RBD binding interface (Figure 4F). Collectively, our structural and functional analyses demonstrate that these convergent somatic mutations contribute substantially to the exceptional neutralization breadth, while the HCDR3 loop and paired light chain also likely constitute the determinants of broad-spectrum recognition. Nevertheless, these IGHV hypermutations not only strengthen neutralizing activity but may also confer resilience against emerging antigenic variants.

### Artificial intelligence-driven discovery of rare and resilient VH3-53/3-66 bnAbs

The existence of bnAbs such as BD55-1205 underscores the feasibility of identifying rare and resilient bnAbs from individuals convalesced or vaccinated during the early phases of a pandemic. We hypothesized that the breadth of neutralization increases with the accumulation of convergent somatic mutations within hotspot paratope residues. To test this, we deeply profiled the antibody repertoires from a cohort of 85 WT vaccine recipients for rare and resilient VH3-53/3-66-encoded bnAbs. Using the WT S-6P probe, we isolated 14,342 antigen-specific B cells from the 85 donors (Figure S7A) and identified 785 monoclonal antibodies with class 1 features that derived from VH3-53/3-66 germlines across ten distinct B cell subsets (Figure S7B-C). We then developed a deep learning workflow incorporating pre-trained language models with dedicated encoders for antigen and antibody sequences (Figure 5A-C). Antigen features were embedded using ESM2^40^, while antibody heavy and light chains were processed using immunoglobulin-specific RoFormer^41^ models trained on 1.2 billion heavy and 210 million light chain sequences from the OAS database^42^. These features were further integrated using convolutional layers and subsequently processed through stacked fully connected (FC) layers to predict the final binding affinity. The model was fine-tuned on a curated dataset of 3,897 RBD-binding antibodies with variant-specific affinity measurements, enabling accurate and variant-aware binding affinity prediction. This model was applied to screen the 785 VH3-53/3-66 antibodies isolated from vaccinees exposed only to ancestral WT antigen. The top 15 candidates, selected based on high predicted affinity for WT, BA.1, and KP.3.1.1, were prioritized for experimental validation (Figure 5D). All the top 15 mAbs showed potent neutralizing activities for both WT and BA.1 strain, and 7 of which cross-neutralize the recent circulating JN.1 variant (Figure 5E). One lead candidate, R102-9, shows potent neutralization to all the five tested strains (Figure 5E). Further deeper validation reveals that R102-9 effectively neutralized all the 19 tested variants, with a mean IC_50_ of 87.8 ng/mL (Figure 5F). The neutralization potency and breadth of R102-9 was superior than BD55-1205 and comparable with the 15 elite VH3-53/3-66 mAbs (Figures 5F, 3A). R102-9 also potently inhibited authentic SARS-CoV-2 WT, XBB.1.5, XBB.1.16, EG.5 and JN.1 isolates with IC_50_ values ranging from 2.6 ng/mL to 37.7 ng/mL (Figure S7D).

**Figure 5.**
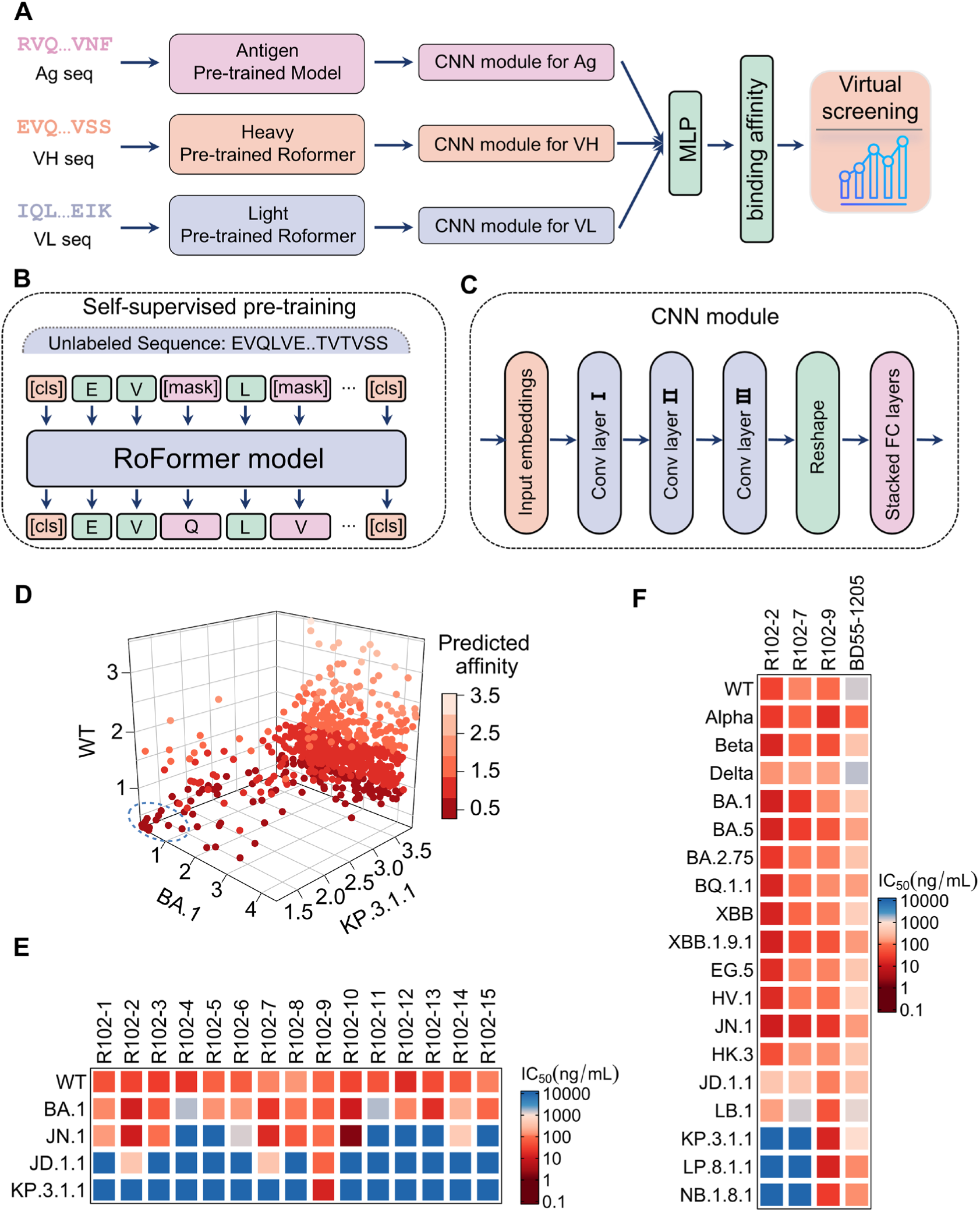
AI-based discovery of VH3-53/3-66 bnAbs. (**A**) Binding affinity prediction and virtual screening were conducted using pre-trained antibody language models and ESM2. The antigen pre-trained model refers to ESM2, which was used to extract evolutionary information from antigen sequences. Heavy and light pre-trained Roformers denote the language models specifically pre-trained on antibody heavy and light chains, respectively, and were employed to capture comprehensive sequence-level features. The extracted features were further processed through dedicated CNN modules for in-depth feature learning, and subsequently used to predict antibody–antigen binding affinity. Based on the affinity prediction model, we scored each antibody in the library and selected those with the highest predicted affinities for wet-lab validation. (**B**) Schematic of self-supervised pre-training. First, tokens in the input sequences were randomly masked, and the RoFormer model was trained to recover the masked regions. In this process, a Unique Amino Acid Tokenizer was employed, which assigns a unique token to each distinct amino acid. (**C**) The CNN module. The input embeddings were processed through three consecutive convolutional layers for in-depth feature extraction, and the final embeddings were generated through stacked fully connected (FC) layers. (**D**) Predicted affinity of 785 VH3-53/3-66 mAbs that bind to WT, BA.1 and KP.3.1.1. The blue dashed line highlights the top 15 antibodies. (**E**) Heatmap showing the neutralizing activity of top15 mAbs against WT, BA.1, JN.1, JD.1.1 and KP.3.1.1. (**F**) Heatmap showing the neutralizing activity of R102-2, R102-7, R102-9, and BD55-1205 against 19 major SARS-CoV-2 strains.

### Accumulated SHMs conferred broad breadth and potency to R102-9

Genetic analysis indicates that R102-9 is encoded by VH3-66, JH4, VK3-15 and JK1, and accumulates ten SHMs in the variable region of the VH3-66 heavy chain. Six of these mutations (F27I^H^, T28I^H^, S31R^H^, V50L^H^, S53P^H^, and Y58F^H^) correspond to previously identified convergent SHMs (Figure 6A), suggesting that our AI modeling is able to recapitulate key SHM features of broadly neutralizing VH3-53/3-66 monoclonal antibodies. Interestingly, introduction of a single G26E^H^ mutation into R102-9 further enhances its neutralization potency (Figure 6B). Notably, the somatically engineered variant, designated R102-9(G26E), exhibits improved potency and breadth compared to both SA55^3^ and Vir-7229^43^ (Figure 6C). To structurally understand R102-9’s exceptional breadth, we determined a cryo-EM structure of the Omicron JD.1.1 S-trimer (S-6P) in complex with R102-9 Fab and C092 Fab, another R1-32-like antibody we previously studied^7^ (Figures 6D, S8-9). Similar to the structure of Omicron LP.8.1 S-trimer bound to ZL58 Fab, we observed a 1:3:3 S-trimer:R102-9:C092 complex and resolved the antibody bound RBD to 2.97 Å resolution through local refinement (Figure 6D, S8-9). Our structure reveals that R102-9 recognizes an epitope nearly identical to that of ZL58, engaging in an extensive network of hydrogen bonds and hydrophobic interactions, with multiple key contacts contributed by the six SHM-derived, predominantly hydrophobic, residues including F27I, T28I, S31R, V50L, S53P, and Y58F (Figure 6E). Taken together, our findings confirm that the predictive power of AI platform is able to identify rare, broadly neutralizing antibodies with exceptional resilience to viral escape from a large collection of mAbs isolated from donors exposed to only WT antigens early in pandemic. These findings demonstrate that, when properly trained, the AI platform can recognize weakened convergent SHM signatures within the noisy immune repertoires of early-pandemic donors to identify rare broadly neutralizing antibodies.

**Figure 6.**
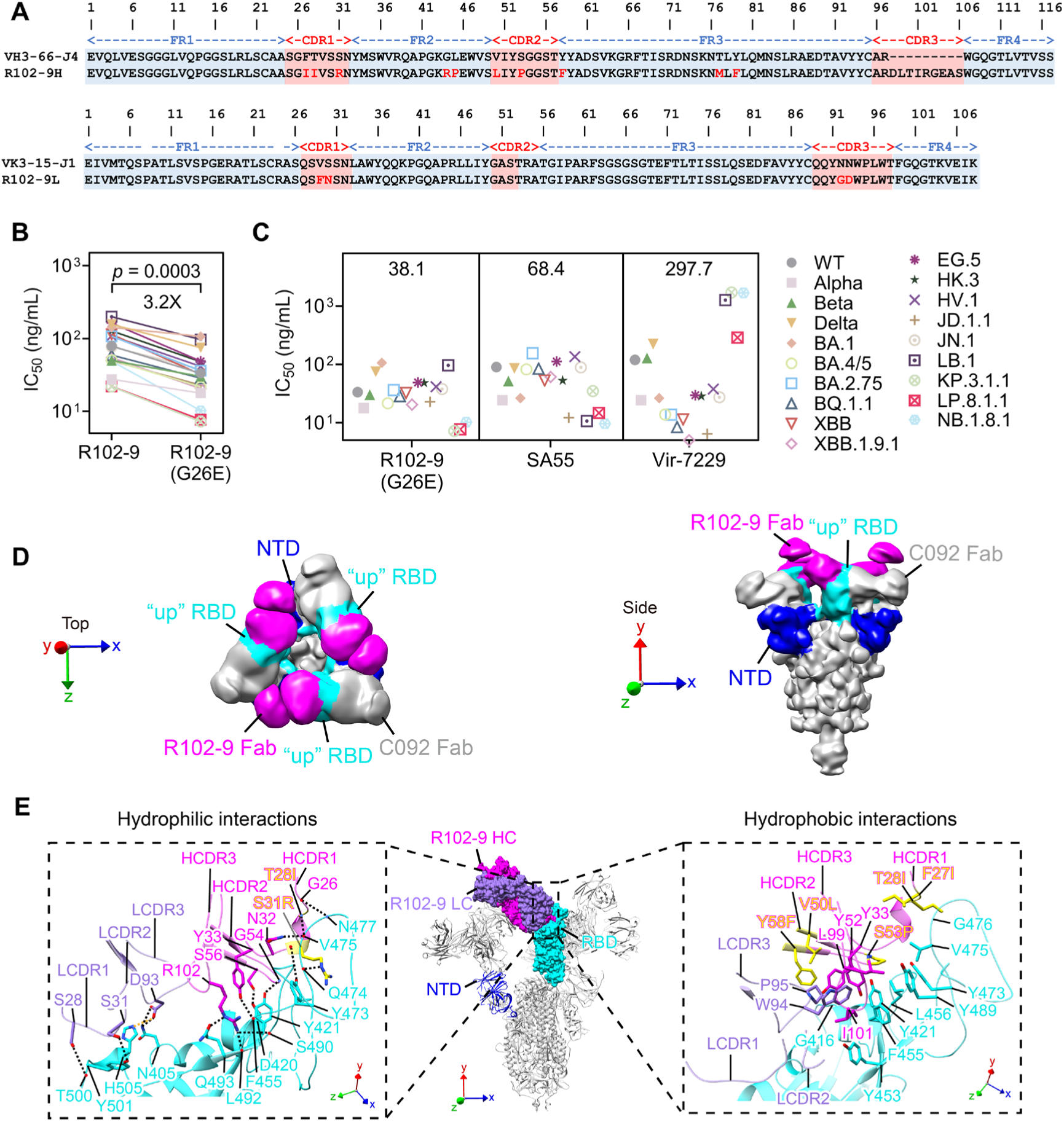
Structural basis of the broad neutralization of R102-9. (**A**) Sequence analysis of R102-9. Somatic hypermutations are highlighted with red. (**B**) Comparison of neutralization activity IC_50_ values between R102-9 and R102-9(G26E). (**C**) Comparison of neutralization activity IC_50_ values across R102-9(G26E), SA55, and Vir-7229. GEM of IC_50_ values were labeled on top. (**D**) Structures (low-pass filtered to 12 Å) of Omicron JD.1.1 spike (S-6P) in complex with R102-9 Fab and C092 Fab. R102-9 Fab, NTD and RBD are highlighted in magenta, blue and cyan, respectively; the rest of the spike and C092 Fab are colored gray. (**E**) Epitope of R102-9 on the RBD of Omicron JD.1.1 spike. Detailed interactions between R102-9 and RBD are shown in the dashed boxes. CDR loops are indicated, and selected interacting residues in antibody-RBD interface are shown. Somatically hypermutated residues are highlighted in yellow and labeled with the changes indicated. Backbone carbonyl oxygens and amide nitrogens are indicated by red and blue dots, respectively. Yellow and black dashed lines indicate salt bridge and hydrogen bond interactions, respectively.

### R102-9 protects from SARS-CoV-2 Beta and JN.1 infection

We next evaluated whether R102-9 protects against SARS-CoV-2 infection in vivo, using BALB/c mice challenged with the Beta variant, which carries the K417N substitution highly detrimental to germline VH3-53/3-66 antibodies. BALB/c mice permit Beta infection with a higher viral load and severer pulmonary pathology in the lungs compared with those infected with Alpha, Delta, and Omicron^44,45^. In both prophylaxis and therapy study, each mouse received a single intraperitoneal injection of either 10 mg/kg high dose, 5 mg/kg intermediate dose or 2 mg/kg low dose of R102-9 (Figure 7A), respectively. Another two group of mice was injected with a positive control SA55^3^ and negative control Influenza-specific mAb MEDI8852^46^ at a high dose of 10 mg/kg. 12 hours before or 24 hours post the mAbs injection, each animal was challenged intranasally with 5×10^4^ focus-forming units (FFUs) of live SARS-CoV-2 Beta strain. The mice were sacrificed for analysis at 3 days post-infection (dpi) when high viral loads and acute lung injury were consistently observed in the negative control group (Figure 7A). In both therapeutic and prophylactic studies, R102-9 conferred robust protection at all tested doses (high, intermediate, and low). Notably, viral titers, determined by FRNT, were undetectable in the lungs of all R102-9-treated mice, whereas animals receiving MEDI8852 exhibited measurable pulmonary viral loads (Figure 7B-C). These findings demonstrate R102-9’s superior efficacy in suppressing Beta variant replication within murine lung tissues. Next, to understand whether or not R102-9 protects against infection-induced lung injury in BALB/c mice, we performed pathological analysis on lung specimens. Compared with MEDI8852-treated mice, mice treated with high or intermediate dose of R102-9 showed mild interstitial alveoli inflammation with minor septal infiltration and congestion (Figure 7D-E). There was limited apparent peribronchiolar infiltration, and the bronchiolar epithelia appeared normal. Mice treated with the low dose showed limited areas of bronchiolar epithelial cell swelling and detachment (Figure 7D-E) together with mild peribronchiolar infiltration and mild alveolar septal infiltration. In contrast, control mice showed large patchy areas of alveolar wall and alveolar space involvement by inflammatory infiltrates and exudation (Figure 7D-E).

**Figure 7.**
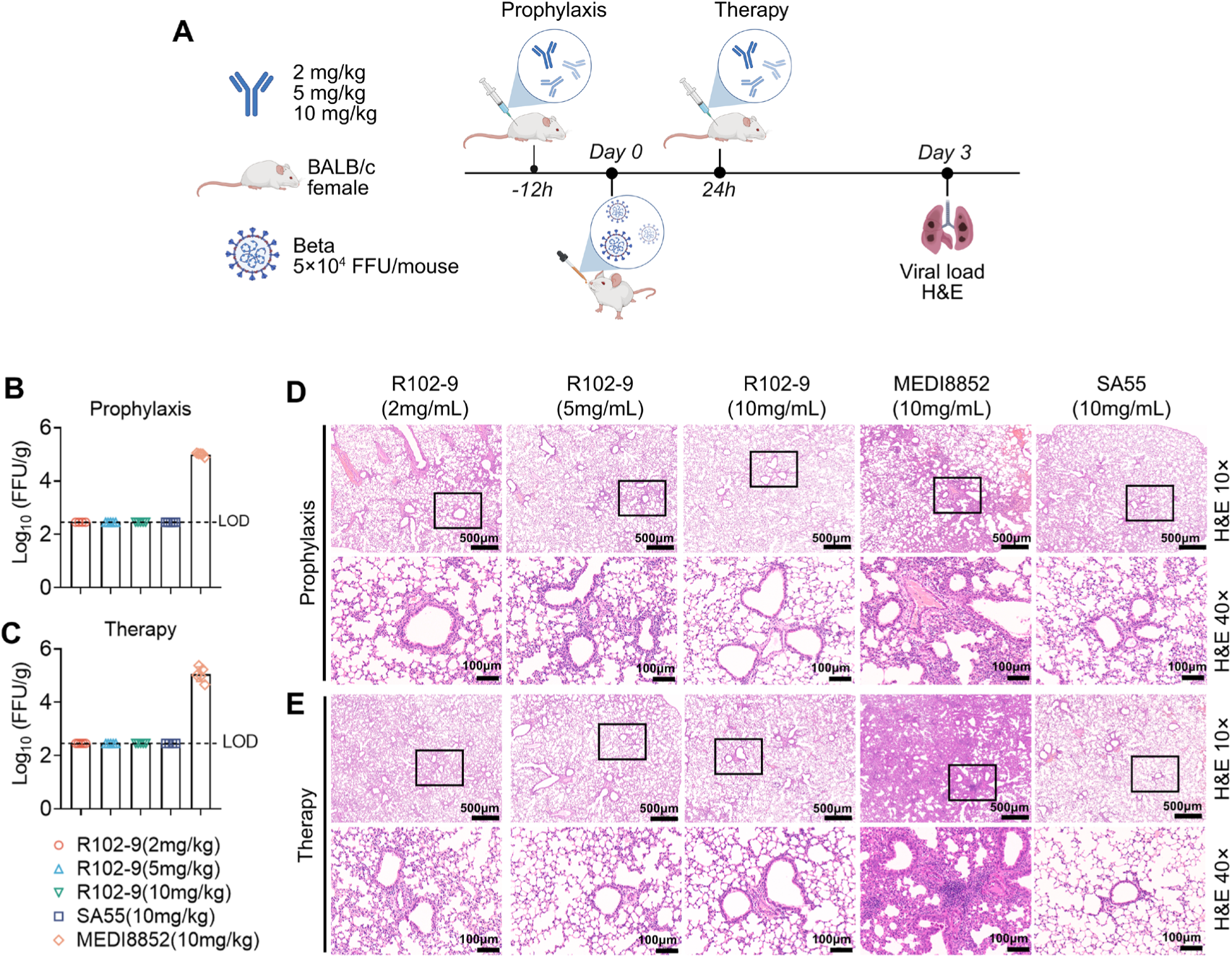
The prophylactic and therapeutic effects of R102-9 against Beta. (**A**) Flowchart of the experimental design of testing the prophylactic and therapeutic activity of R102-9 to protect against challenge with Beta SARS-CoV-2 in BALB/c mice. (**B-C**) Infectious virions were tested by viral fluorescent assay in lung tissue homogenates of prophylactic (**B**) and therapeutic (**C**) assay. FFUs per g of tissue extractions were compared between different groups in log_10_-transformed units. LOD, limit of detection. (**D-E**) Representative images of mouse lung tissues of prophylactic (**D**) and therapeutic (**E**) by H&E staining.

To further evaluate the protective efficacy of R102-9 against emerging variants, we challenged K18-hACE2 mice with 10,000 FFU of the JN.1 variant (Figure S10A). Mice received a low dose of R102-9 (2 mg/kg) either 12 hours before or after infection. MEDI8852 and SA55 were included as positive and negative controls, respectively. Both prophylactic and therapeutic administration of R102-9 markedly reduced pulmonary viral titers (Figure S10B-C), and histopathological analysis revealed attenuated lung inflammation and improved tissue histological features in treated animals (Figure S10D-E). These results demonstrate that the AI-discovered antibody R102-9 exhibits robust protective efficacy in animal model.

## Discussion

This study delineates the molecular evolution of durable, broadly neutralizing VH3-53/3-66 antibodies in elite neutralizers with sequential Omicron breakthrough infections. Repeated antigenic exposures drive convergent somatic hypermutations that markedly expand breadth and potency, preserving neutralization even against the most recent immune-evasive variants (e.g., JD.1.1, LP.8.1.1, NB.1.8.1). Our findings extend prior reports that VH3-53/3-66 antibodies are public clonotypes^11,47,48^ by showing how recurrent antigenic exposures sculpt their affinity maturation into pan-variant neutralizers. Overall, despite the high variability of the VH3-53/3-66–targeted epitope, we show that the immune system can still generate broad and potent neutralizing antibodies directed against this epitope.

Previous studies showed that VH3-53/3-66 antibodies isolated early in the pandemic (e.g., CB6, P2C-1F11) had limited breadth owing to minimal SHMs, showing vulnerability to K417N/T substitutions^49^. Although convergent SHMs that improved affinity and breadth against early Omicron variants were reported^26–28,50^, these antibodies fail to effectively neutralize more recent escape mutants (Figure 3A). In contrast, our analysis indicates that sequential Omicron exposures drive accumulation of additional key SHMs (G26E, F27I, T28I, S31R, V50L, S53P, Y58F) that collectively strengthen paratope-RBD interactions. Structurally, ZL58 and R102-9 illustrate how these SHMs introduce compensatory contacts at escaped epitopes, paralleling VacBB-551^23^ and 19-77^51^ yet exhibiting greater resilience to A475V/L455F/F456L in JD.1.1 and LP.8.1.1. These results reveal two layers of epistasis: mutations within SARS-CoV-2 spike epitopes show strong genetic interactions ^52^, and the SHMs accrued during affinity maturation of VH3-53/3-66 broadly neutralizing antibodies likewise follow a cooperative, epistatic pattern. These bidirectional genetic interactions provide compelling evidence for co-evolutionary dynamics between viral antigens and neutralizing antibodies.

Critically, we show that germline-reverted antibodies lose breadth, while introducing these convergent SHMs into clinically escaped antibodies (BD-604, P2C-1F11, LY-CoV488) restores neutralization against Omicron lineages (380-fold potency increase for BD-604). This strategy diverges from conventional engineering approaches by leveraging natural selection patterns^51^. Restoring the activity of escaped antibodies through SHM grafting provides a rapid strategy to update clinical antibody candidates against emerging variants. For vaccines, our data suggest that iterative Omicron exposures, akin to prime-boost regimens with divergent spikes^53^, foster bnAb maturation. Notably, mucosal IgA^+^ VH3-53/3-66 clones were enriched in elite neutralizers, hinting at synergistic systemic/mucosal responses, an underexplored area in current vaccine design^54^.

While most VH3-53/3-66 antibodies isolated early in the pandemic lack breadth, rare exceptions such as BD55-1205 suggest that some clones were pre-adapted to future variants^19^. Our AI-guided discovery of R102-9 advances bnAb identification by showing that machine learning can extract critical SHM signatures from early repertoires, a capability not achieved previously^19,55^. R102-9 neutralizes all 19 tested variants (mean IC_50_: 87.8 ng/mL) and protects mice against Beta and JN.1 challenge, outperforming BD55-1205 and matching elite antibodies induced by sequential Omicron breakthrough infections. This approach diverges from prior epitope-focused AI tools^56,57^ by linking SHM patterns to function, enabling rapid mining of resilient bnAbs without extensive laborious experimental screening^58^. Structurally, R102-9’s mode of action parallels ZL58, underscoring SHM signatures as predictive features of breadth. This study establishes a new workflow for pandemic preparedness: mining early immune repertoires prospectively for rare, resilient bnAbs rather than waiting for elite neutralizers to emerge.

In summary, we chart a molecular roadmap for the evolution of broadly protective VH3-53/3-66 antibodies under sequential exposures, identifying seven key convergent SHMs that collectively overcome SARS-CoV-2 immune evasion. Integrating repertoire analysis, antibody engineering, structural biology, and AI, we show that these naturally selected patterns can restore escaped therapeutic antibodies and uncover rare, potent bnAbs from early-pandemic repertoires. This work elucidates the coevolution between a public antibody class and the viral surface antigen and offers actionable guidance for variant-resilient countermeasures, establishing a practical, machine learning–enabled antibody discovery workflow for pandemic preparedness.

### Limitations of the study

While this study provides key insights into the evolution of broad SARS-CoV-2 neutralization, several limitations should be noted. First, due to the limited number of B cells that could be sorted from each individual, our B cell sequencing was performed on pooled samples rather than analyzing cells from each donor separately. This approach may obscure individual-specific immune responses and limit our ability to track clonal evolution at the single-donor level. Second, current analytical constraints prevented precise identification of somatic hypermutations in the CDR3 region. As a result, our study primarily focused on SHM patterns in CDR1 and CDR2, which may not fully capture the mutational landscape contributing to antibody breadth and potency. Third, the AI-based prediction framework, while promising, was specifically trained and validated for the VH3-53/3-66 antibody lineage, expansion to other antibody classes (e.g., VH1-69 or VH1-58) will be necessary to determine its generalizability. These limitations highlight important directions for future research, including single-donor sequencing approaches, improved CDR3 mutation analysis methods, and development of lineage-agnostic prediction models. Fourth, the SHM convergence signal is relatively weak among antibody sequences isolated early in the pandemic, and obtaining sufficiently large datasets for a given clonal type may be challenging.

## Resource availability

### Lead contact

Further information, requests for resources, and reagents should be directed to and will be fulfilled by the lead contact, Jincun Zhao.

### Materials availability

All expression plasmids and proteins generated in this study are available upon request.

### Data and code availability

Single-cell RNA-seq raw data sequencing data have been deposited at National Genomics Data Center and are available under the accession BioProject ID PRJCA049739. Cryo-EM density maps for the SARS-CoV-2 Omicron LP.8.1 spike in complex with ZL58 and ZL525 Fabs, and the Omicron JD.1.1 spike in complex with R102-9 and C092 Fabs have been deposited in the Electron Microscopy DataBank (EMDB) with accession codes EMD-66901, EMD-66902, EMD-66903, and EMD-66904 respectively. The related atomic models have been deposited in Protein Data Bank (PDB) under accession codes 9XI9, 9XIB, 9XIC, and 9XID respectively. All maps and models are publicly available as of the date of publication. Accession numbers are listed in the key resources table. The code used in AI-based antibody discovery is available at: https://github.com/YidongSong/Ab-Ag-AFF/tree/main. Any additional information required to reanalyze the data reported in this work paper is available from the lead contact upon request.

## Methods

### Cells and viruses

Expi293F cells (Thermo Fisher Scientific, A14527) were maintained in Expi293F Expression medium (Thermo Fisher Scientific, A1435101) at 37 °C by shaking at 120 rpm under a humidified atmosphere with 8% CO_2_. Human embryonic kidney (HEK) 293T (ATCC, Cat# CRL-3216), HEK293T-hACE2, Vero E6 (ATCC, Cat# CRL-1586) and Huh-7 cells (JCRB, Cat# 0403) were maintained in 10% FBS supplemented Dulbecco’s Modified Eagle’s Medium (DMEM) at 37°C, 5% CO_2_. The authentic SARS-CoV-2 viruses, including WT, BA.5, XBB.1, XBB.1.5, XBB.1.16, EG.5, and JN.1 were isolated from COVID-19 patients and preserved in Guangzhou Customs District Technology Center BSL-3 Laboratory. Experiments related to authentic SARS-CoV-2 viruses were conducted in Guangzhou Customs District Technology Center BSL-3 Laboratory.

### COVID-19 patient and vaccine recipient enrollment

A cohort of 68 individuals with confirmed SARS-CoV-2 infection was enrolled between December 2022 and December 2023. All participants experienced sequential breakthrough infections with the BA.5 and XBB subvariants during the two major epidemic waves in this period. Each individual had previously received three doses of an inactivated COVID-19 vaccine (Sinopharm/CoronaVac). SARS-CoV-2 infection was confirmed by RT–PCR. An additional cohort of 85 vaccine recipients (WT-vaccinated group) was included. This group comprised 43 individuals who received three doses of an inactivated vaccine (Sinopharm/CoronaVac), 11 who received two doses of an inactivated vaccine (Sinopharm/CoronaVac) followed by one adenoviral vector vaccine (CanSino), 18 who received two doses of an inactivated vaccine (Sinopharm/CoronaVac) followed by one mRNA vaccine (BioNTech), and 13 who received three doses of an mRNA vaccine (BioNTech). All participants provided written informed consent prior to enrollment. The study was conducted under the approval and supervision of the Ethics Committee of the First Affiliated Hospital of Guangzhou Medical University (Ethics IDs: ES-2020-65, ES-2023-015-01, ES-2021-78).

### Protein expression and purification

The gene encoding the SARS-CoV-2 spike (S) protein extracellular domain (residues 14–1211) was cloned into the pcDNA3.1(+) mammalian expression vector. The construct included an N-terminal mu-phosphatase signal peptide, an “R” substitution at the furin cleavage site (residues 682–685), a C-terminal TEV protease cleavage site, a T4 fibritin trimerization motif, and a His₆ tag, designated “S-R”, as previously described^59^. For structural studies, a stabilized S-protein variant (“S-GSAS/6P”) carrying six proline substitutions (F817P, A892P, A899P, A942P, K986P, and V987P) was engineered and termed “S-6P”^60^. The RBD (residues 319–541) was constructed with an N-terminal mu-phosphatase signal peptide and a C-terminal Avi-tag followed by a 6×His tag, and similarly cloned into pcDNA3.1(+). All constructs were transiently transfected into Expi293F cells using polyethylenimine. Proteins were purified by immobilized metal affinity chromatography (IMAC) under previously established conditions^7,61^, aliquoted, flash-frozen in liquid nitrogen, and stored at −80°C.

### B cell sorting and single-cell RNA sequencing

B cells were enriched from peripheral blood mononuclear cells (PBMCs) using an EasySep Pan-B Cell Magnetic Enrichment Kit (STEMCELL Technologies) and incubated with 2 mg/mL of Avi-tagged JD.1.1 or KP.3.1.1 RBD on ice for 30 minutes. Cells were subsequently washed with PBS containing 2% fetal bovine serum (FBS; Hyclone) and stained with a cocktail of anti-human CD3 (PerCP-Cy5.5; BioLegend, Cat# 410710) and anti-human CD19 (FITC; BioLegend, Cat# 348205) antibodies. To identify RBD-specific B cells, PE- and APC-conjugated anti-Avi-tag antibodies (BioLegend, Cat# 637309 and #637307) were included, enabling the gating of double-positive cells. RBD-specific double-positive B cells were gated as DRAQ7⁻CD3⁻CD19⁺ and sorted on a FACSAria II cell sorter. Sorted B cells were processed using the following 10x Genomics kits: Chromium Next GEM Single Cell 5′ Kit v3, Library Construction Kit, Chromium Next GEM Chip K Single Cell Kit, Chromium Single Cell Human BCR Amplification Kit, and Dual Index Kit TT Set A. Following GEM generation and barcoding, cDNA was synthesized via GEM reverse transcription (RT) and purified using bead-based cleanup. The purified cDNA was amplified for 10–14 cycles and cleaned with SPRIselect beads. cDNA quality and concentration were assessed using a 4200 TapeStation System (Agilent Technologies). BCR enrichment was performed on full-length cDNA. Libraries for gene expression and BCR analysis were prepared according to the Chromium Next GEM Single Cell 5′ Reagent Kits v3 User Guide, with PCR cycle numbers adjusted based on cDNA concentration. Final libraries were sequenced on a NovaSeq S4 instrument (Illumina), with a target median depth of 50,000 read pairs per cell for gene expression libraries and 5,000 read pairs per cell for BCR libraries.

### Single-cell RNA-seq analysis

Single-cell RNA sequencing and BCR sequencing data were processed with Cell Ranger (v7.0.0) and aligned to the GRCh38-2020 human reference genome provided by 10x Genomics. Count matrices were analyzed in R (v4.0.2) using the Seurat package (v5.2.1). Cells were filtered based on the following criteria: mitochondrial gene content < 20% and feature count < 4,000. The three sequencing runs were integrated using log-normalized counts and a canonical correlation analysis approach with 2,000 highly variable genes. The combined object was subjected to principal component analysis (PCA); the top 30 principal components were used for Uniform Manifold Approximation and Projection (UMAP) embedding and neighbor identification. Clustering was performed at a resolution of 0.6. For BCR analysis, filtered contig annotations from Cell Ranger V(D)J were imported into R and processed using the scRepertoire package (v1.1.3). For mutation analysis, heavy and light chains from monoclonal antibodies and single-cell BCRs were annotated with IgBLAST (v1.14.0) using IMGT human reference genes. Resulting files were parsed with Change-O (v0.4.6). Mutation frequency was computed as previously described^62^ using the *calcObservedMutations* function from SHazaM (v1.0.2), by counting nucleotide and amino acid mismatches relative to the germline sequence within the variable region up to the CDR3.

### Expression of monoclonal antibody

Heavy- and light-chain variable region genes (VH and VL) were cloned into human IgG1 expression vectors using the CloneExpress II One-Step Cloning Kit (Vazyme, Cat# C112-02). HEK293F cells were cultured to a density of 1 × 10⁶ cells/mL and co-transfected with equimolar amounts of heavy- and light-chain plasmids using the EZ Cell Transfection Reagent (Life-iLab Biotech, Cat# AC04L092). Transfected cells were maintained in CD 293 TGE medium (ACRO, Cat# CM-1156) supplemented with 10% CD Feed X (ACRO, Cat# CF-1116-12) at 37 °C under 5% CO₂ with shaking at 120 rpm. Six days post-transfection, culture supernatants were harvested, clarified by centrifugation, and filtered through 0.22-μm membranes (Merck Millipore, Cat# SLGP033NS). Filtered supernatants were incubated with Protein A Resin (Genscript, Cat# L00210) for 2 hours at room temperature for affinity purification. After extensive washing, bound antibodies were eluted with 0.1 M sodium citrate (pH 3.25) and immediately neutralized with 1 M Tris-HCl (pH 8.8). Purified antibodies were concentrated in PBS using Amicon Ultra centrifugal filters (Merck Millipore, Cat# UFC810096) and stored at −80 °C.

### Pseudovirus production and neutralization assay

The pseudovirus was constructed based on a VSV-G pseudotyped backbone in which the native G gene was replaced by a firefly luciferase (Fluc) reporter gene and the SARS-CoV-2 spike protein was incorporated. The SARS-CoV-2 WT and variant spike genes were codon-optimized and cloned into the pcDNA3.1 vector to generate the expression plasmid pcDNA3.1-SARS-CoV-2-spikes. HEK293T cells were trypsinized, resuspended at a concentration of 5–7 × 10⁵ cells/mL, and seeded into a T75 flask with 15 mL of cell suspension. After overnight incubation at 37 °C under 5% CO₂, cells reached 70–90% confluence. The medium was then removed, and cells were infected with 15 mL of G*ΔG-VSV pseudotyped virus (Kerafast) at a titer of 7.0 × 10⁴ TCID₅₀/mL. Concurrently, 30 μg of the spike expression plasmid was transfected using Lipofectamine 3000 (Invitrogen, L3000015) according to the manufacturer’s instructions, followed by 6–8 h of incubation at 37 °C with 5% CO_2_. After transfection, the supernatant was aspirated, and cells were gently washed twice with PBS containing 1% FBS. Fresh complete DMEM (15 mL) was added, and cells were cultured for 24 h. The supernatant was collected, and another 15 mL of fresh complete DMEM was added for a further 24 h incubation. The two batches of supernatant were combined, centrifuged at 1000 × g for 10 min, filtered through a 0.45-μm membrane, aliquoted, and stored at −80 °C for subsequent use. For pseudovirus neutralization assays, Huh-7 cells were employed. Test antibodies were serially diluted six times in complete DMEM in 96-well plates. Pseudovirus was added at a concentration of 1.3 × 10⁴ TCID₅₀/mL per well, and the mixture was incubated for 1 h at 37 °C under 5% CO₂. Then, 100 μL of trypsinized Huh-7 cells (2 × 10⁵ cells/mL, 0.25% Trypsin-EDTA, Gibco) were added to each well. After 24 h of incubation, the supernatant was carefully removed leaving 100 μL per well, and 100 μL of luciferase substrate (PerkinElmer, 6066769) was added. Following a 2-minute incubation at room temperature, 150 μL of the lysate was transferred to a white opaque 96-well microplate (PerkinElmer), and luminescence was measured using an Ensight multimode plate reader (PerkinElmer). All samples were assayed in duplicate. The half-maximal inhibitory concentration (IC₅₀) was determined by four-parameter nonlinear regression analysis using GraphPad Prism version 8.0.

### SARS-CoV-2 authentic virus neutralization assay

Antibodies were serially diluted in DMEM and mixed with 200 focus-forming units (FFU) of authentic SARS-CoV-2 viruses, including WT, BA.5, XBB.1, XBB.1.5, XBB.1.16, EG.5, and JN.1. Following incubation at 37 °C for 1 h, the antibody–virus mixtures were applied to Vero E6 cells cultured in 96-well plates and incubated for 1 h at 37 °C under 5% CO₂. After removal of the inoculum, each well was overlaid with 100 μL of pre-warmed 1.6% carboxymethylcellulose and cultured for 24 h. The overlay was then aspirated, and cells were fixed with 4% paraformaldehyde (Biosharp, Cat# BL539A) and permeabilized with 0.2% Triton X-100 (Sigma, Cat# T8787). Subsequently, cells were incubated for 1 h at 37 °C with a human anti-SARS-CoV-2 nucleocapsid protein monoclonal antibody (generated by in-house screening). After three washes with 0.15% PBST, an HRP-conjugated goat anti-human secondary antibody (Jackson ImmunoResearch, Cat# 609-035-213) was applied and incubated for 1 h at 37 °C. Following three additional washes with 0.15% PBST, foci were developed using TrueBlue Peroxidase Substrate (KPL, Cat# 50-78-02) and quantified with an ELISPOT reader (Cellular Technology Ltd.). The 50% foci reduction neutralization titer (FRNT_50_) was determined by four-parameter nonlinear regression analysis in GraphPad Prism 8.0.

### Binding affinity prediction, Virtual screening of the VH3-53/3-66 antibody library

To predict antibody–antigen binding affinities, we employed pre-trained language models tailored to the distinct properties of antigens and antibodies (Figure 5A). Antigen sequences were encoded using ESM2^40^ to capture conserved evolutionary features. For antibody sequences, we developed specialized language models trained on immunoglobulin-specific data. As illustrated in Figure 5B, we preprocessed unlabeled antibody sequences using a Unique Amino Acid (UAA) Tokenizer, and pre-trained the RoFormer model^41^ on 1.2 billion heavy chain and 210 million light chain sequences from the OAS database^42^ via masked language modeling, yielding separate pre-trained models for heavy and light chains. Features were extracted from multiple chains using pre-trained models, and binding affinity was predicted through the convolutional neural network (CNN) module comprising consecutive convolutional layers and stacked fully connected (FC) layers (Figure 5C). The model was fine-tuned on a custom-curated dataset integrating in-house and public sources^16,18,52,63–68^, comprising 3,897 RBD-binding antibodies with quantitative affinity measurements across multiple SARS-CoV-2 variants, including WT, Beta, Delta, BA.1, BA.5, and the recently emerged KP.3.1.1 strain. This enabled the model to learn variant-specific binding profiles with high accuracy. We then applied this optimized framework to virtually screen a library of 785 VH3-53/3-66-encoded monoclonal antibodies. Each antibody was computationally paired with target antigens, and binding affinities were predicted to rank candidates. Iterative screening across variants enabled identification of antibodies with strong and broad neutralizing potential.

### Cryo-EM sample preparation and data collection

For the LP.8.1 S-6P:ZL58 Fab:ZL525 Fab complex, LP.8.1 S-6P at 5.1 mg/ml was mixed with ZL58 Fab and ZL525 Fab at 1:1:1 molar ratio. For the JD.1.1 S-6P:R102-9 Fab:C092 Fab complex, JD.1.1 S-6P at 6 mg/ml was mixed with R102-9 Fab and C092 Fab at 1:1:1 molar ratio. After incubation for 1 min at room temperature, 3 µL of each mixture supplemented with 0.1% octyl-glucoside (Sigma-Aldrich, Cat# V900365) was applied to freshly glow-discharged (30 s at 15 mA) holey carbon grids (Quantifoil, Cu R1.2/R1.3). The grids were blotted for 2.5 s with a blot force of 4, and then plunge-frozen into liquid ethane using a Vitrobot Mark IV (Thermo Fisher Scientific) at 18 °C and 100% humidity. Cryo-EM grids were loaded onto a 300 keV Titan Krios electron microscope equipped with a Falcon 4 direct electron detector with SelectrisX energy filter (slit width 10 eV). Movie stacks were automatically collected using EPU software with the electron event representation (EER) mode at a nominal magnification of ×160,000 with a pixel size of 0.73 Å and a defocus range from -0.6 to -2.0 µm. Each stack was recorded and exposed at a dose rate of 6.53 e^-^/pixel/s for 4.08 s, resulting in a total dose of ∼50 e^-^/Å^2^.

### Cyro-EM data processing

Data processing was performed using cryoSPARC v4.2.0. After patch motion correction, contrast transfer function (CTF) estimation and removal of poor-quality micrographs, 696,375 and 955,134 particles were picked from 6,117 micrographs of the LP.8.1 S-6P:ZL58 Fab:ZL525 Fab dataset and 6,579 micrographs of the JD.1.1 S-6P:R102-9 Fab:C092 Fab dataset, respectively. Subsequent 2D classification was used to remove low-quality particles, and well-defined particles were selected for ab-initio reconstruction and heterogeneous refinement. For the JD.1.1 S-6P:R102-9 Fab:C092 Fab dataset, additional non-uniform refinement was applied. Particles from classes exhibiting high-resolution features were used to train the topaz picker, followed by particle re-extraction, 2D classification, ab-initio reconstruction, heterogeneous refinement, and non-uniform refinement. This workflow yielded 3:3:3 LP.8.1 S-6P:ZL58 Fab:ZL525 Fab and 3:3:3 JD.1.1 S-6P:R102-9 Fab:C092 Fab complexes at global resolutions of 2.75 Å and 2.76 Å, respectively. To improve the resolution of the RBD-Fab binding interface, a focused mask covering the RBD-Fab region was generated from the globally refined maps. Using this mask, 3D classification was performed to select particles with well-defined local features, followed by local refinement, yielding focused maps at 2.93 Å and 2.97 Å, respectively. Map resolutions were estimated at the 0.143 criterion of the phase-randomization-corrected Fourier shell correlation (FSC) curve calculated between two independently refined half-maps multiplied by a soft-edged solvent mask. The data processing procedures are summarized in Figures S5, S6, S8, and S9. The estimated B-factors of maps are listed in Table S3.

### Model building and structural analysis

A previously determined structure of the SARS-CoV-2 WT S-trimer in complex with 3 YB8-259 Fabs and 3 R1-32 Fabs (PDB: 8HC4) and a structure of the SARS-CoV-2 WT S1 in complex with a YB9-258 Fab and an R1-32 Fab (PDB: 8HC5)^50^ were used as starting models and fitted into final refined maps in UCSF Chimera v1.15^69^. Iterative model building and real space refinement were carried out in Coot v0.9.8.1^70^ and PHENIX v1.19.1^71^. Model refinement statistics are summarized in Table S3. Interface analyses were performed in PISA^72^. Structure figures were generated in UCSF Chimera v1.15.

### SARS-CoV-2 infection and treatment in mice

In vivo prophylactic and therapeutic activities of R102-9 against SARS-CoV-2 Beta variant were assessed in BALB/c mice. Infection studies were performed on 6 to 8 weeks-old female mice, in animal biosafety level 3 (BSL-3) facilities of Guangzhou Customs District Technology Center. Animal work was approved by the Animal Experimentation Ethics Committee of the First Affiliated Hospital of Guangzhou Medical University (IACUC number: 20241225). Briefly, anesthetized mice were inoculated intranasally with 5×10^4^ FFU of SARS-CoV-2 Beta variant. 12 hours before inoculation (prophylactic group) or 24 hours post inoculation (therapeutic group), mice received an intraperitoneal injection with 2, 5, or 10 mg/kg of R102-9. Mice injected with 10mg/kg MEDI8852 or SA55 were challenged with the same dose of SARS-CoV-2 Beta as controls. Lungs were harvested at 3 days post infection for virus titration and hematoxylin and eosin (H&E) staining. In the JN.1 protection experiment, we intranasally challenged K18-hACE2 transgenic mice with 10,000 FFU of the JN.1 variant under light anesthesia. R102-9 antibody was administered intramuscularly at a dose of 2 mg/kg either 12 hours before (prophylactic group) or 12 hours after (therapeutic group) viral challenge.

### Statistical analysis

All statistical analyses were performed using R (v4.4.0) or GraphPad Prism (v8.0). Details of the specific tests used are provided in the corresponding figure legends. Participant characteristics were summarized as counts (n), medians with interquartile ranges (IQR), or frequencies with percentages. Chi-square and Fisher’s exact tests were used to assess differences in categorical variables. Quantitative variables were compared using the Wilcoxon matched-pairs test for within-group analyses and the Mann–Whitney U test for comparisons between independent groups.

## Acknowledgments

We thank the Huimin Feng at the Cryo-EM Facility of GIBH-CAS and Mei Li at Guangzhou Laboratory Bio-imaging Technology Platform for help with cryo-EM sample preparation and data collection. We thank the Drug Discovery Platform of Guangzhou National Laboratory for our ITC experiments, and we would be grateful to Tianyong Tu for his help with sample preparation. This study was supported by the National Natural Science Foundation of China (82025001, 82495200 and 82495203 to J.Z., 82341085 and 32570199 to X.X., and 82201932 to Q.Y.), the R&D Program of Guangzhou Laboratory (EKPG21-30-2 to J.Z. and SRPG22-002 to X.X.), the Basic Research Project of Guangzhou institutes of Biomedicine and health, Chinese Academy of Sciences (GIBHBRP24-02 to X.X.), the Natural Science Fund of Guangdong Province (2025A1515011245 to B.L.), and the Young Doctoral Starting Sail Project of the Guangzhou Municipal Science and Technology Bureau (2024A04J4195 to B.L.).

## Author contributions

J.Z. and Q.Y. conceived and initiated the study; X.X. conceived and initiated the structural studies; X.H., Y.Y., H.Z, Z.H., and S.L. performed antibody isolation, expressed, and purified RBDs and antibodies; X.H., Y.Y., H.Z, Z.H., F.C.,Y.L., L.C., N.Z., J.X., C.C., and H.B. performed the pseudovirus neutralization assays; H.X. performed authentic virus neutralization assays and animal challenge assays under the supervision of A.Z.; Q.Y., T. W., J.L, and M.W. performed the bioinformatic analysis; B.L., C.N., and Z.L. collected cryo-EM data under the supervision of X.X.; B.L., C.N., and Z.L processed cryo-EM data under the supervision of X.X.; B.L. C.N., Z.L., H.Y., and X.X. analyzed cryo-EM structures; Y.S. and F.W. performed the AI-based antibody prediction under the supervision of J.Y. and YD.Y.; Y.Q., X.H., B.L., Y.Y., H.Z., and Y. S. prepared the figures; Q.Y., B.L., and Y.S. wrote the paper with input from all authors. JX.Z., J.Y., YD.Y., A.Z., X.X., and J.Z. supervised the research.

## Supplementary figures

**Figure S1.**
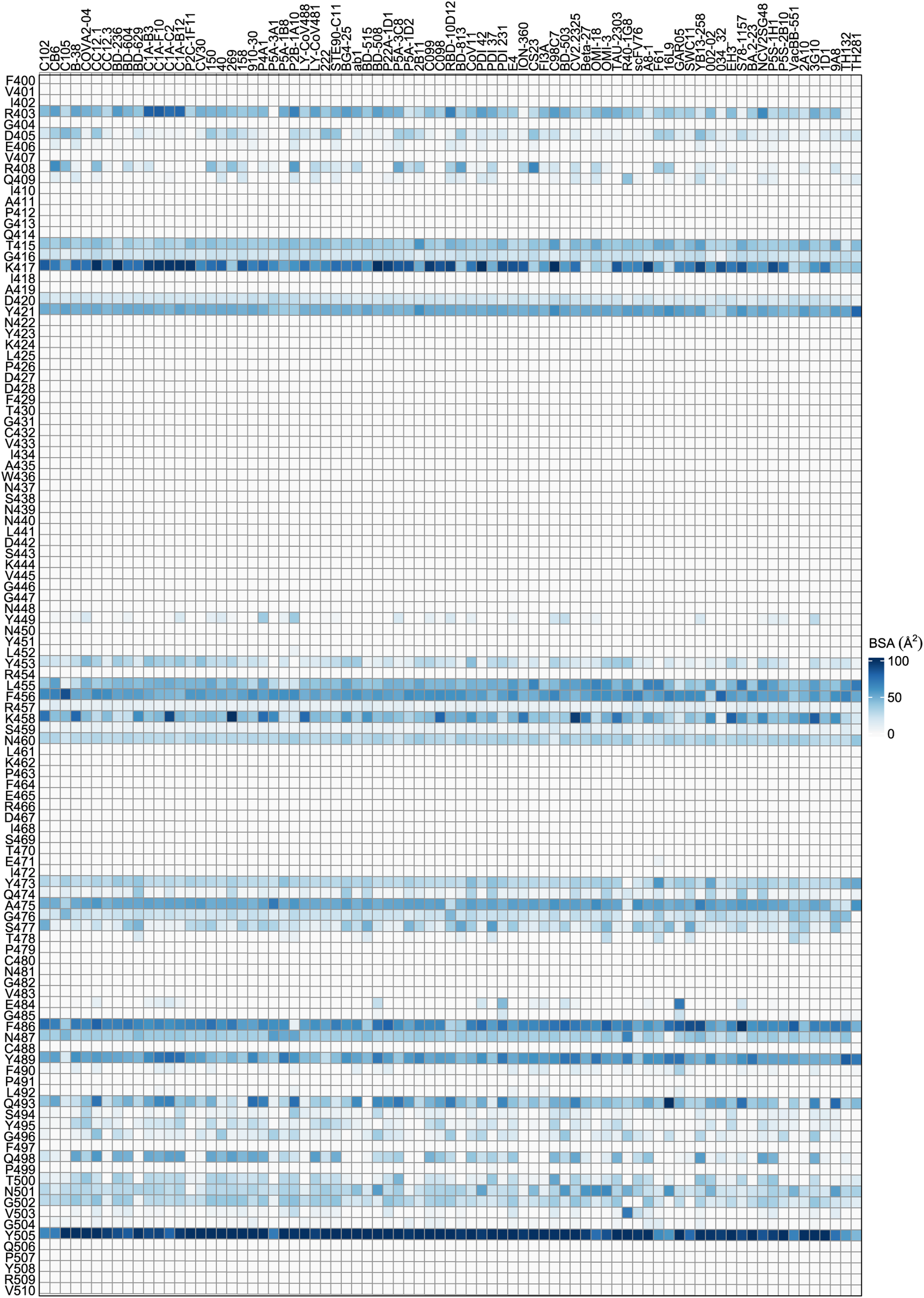
Heatmap showing the buried surface area of 79 class 1 VH3-53/3-66 mAbs at each epitope sites on RBD.

**Figure S2.**
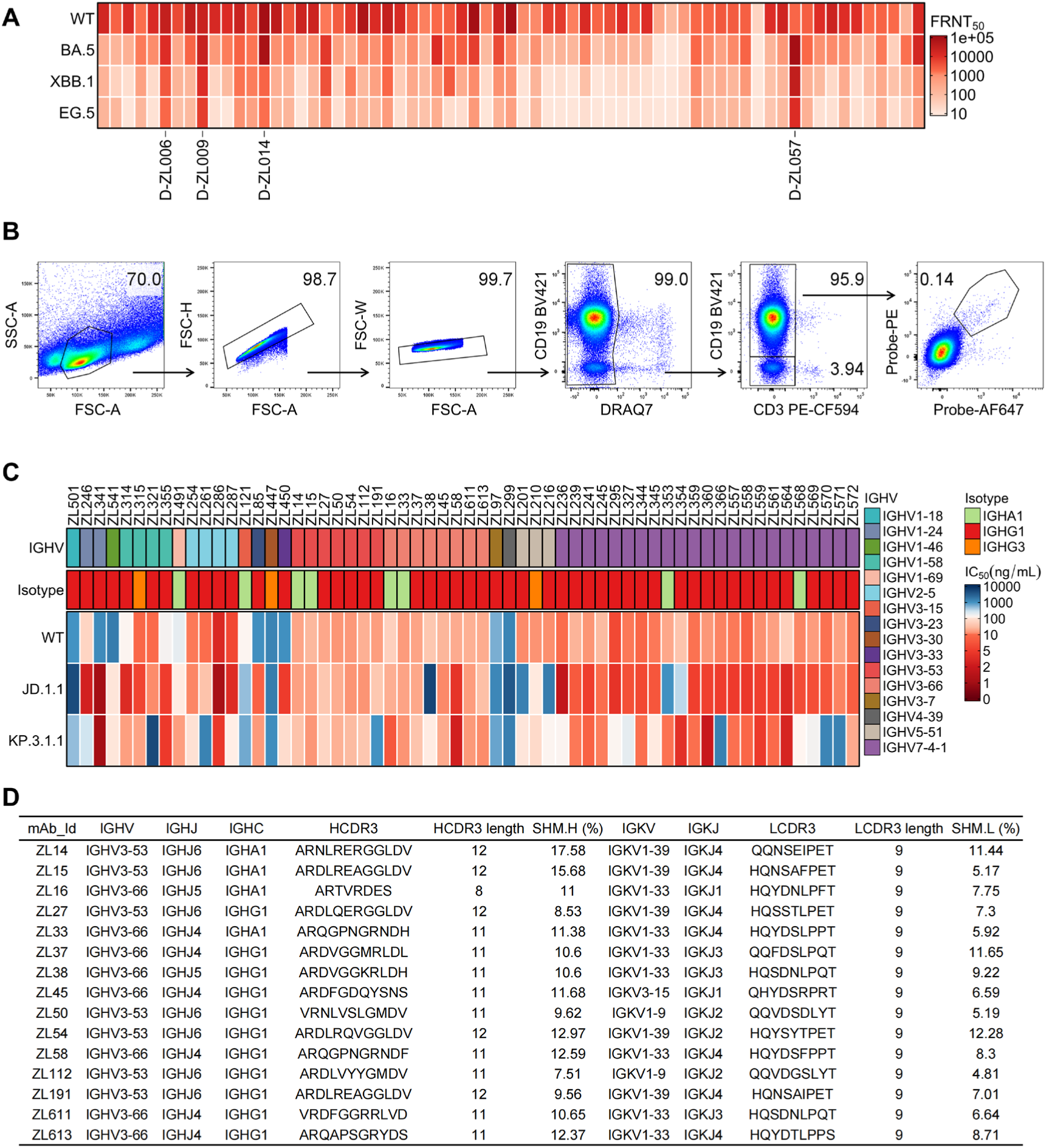
Isolation of broad neutralizing mAbs from elite neutralizers. (**A**) Heatmap showing the neutralizing activity of serum of 68 individuals with BA.5 and XBB breakthrough infections. (**B**) The flow cytometry gating strategy for sorting antigen-specific B cells. (**C**) Heatmap showing the IC_50_ values of 60 cross-neutralizing mAbs against WT, JD.1.1 and KP.3.1.1. The top annotations indicate the IGHV and IGHC (isotype) germline segments. (**D**) The summary of genetic characters of the 15 VH3-53/3-66 mAbs.

**Figure S3.**
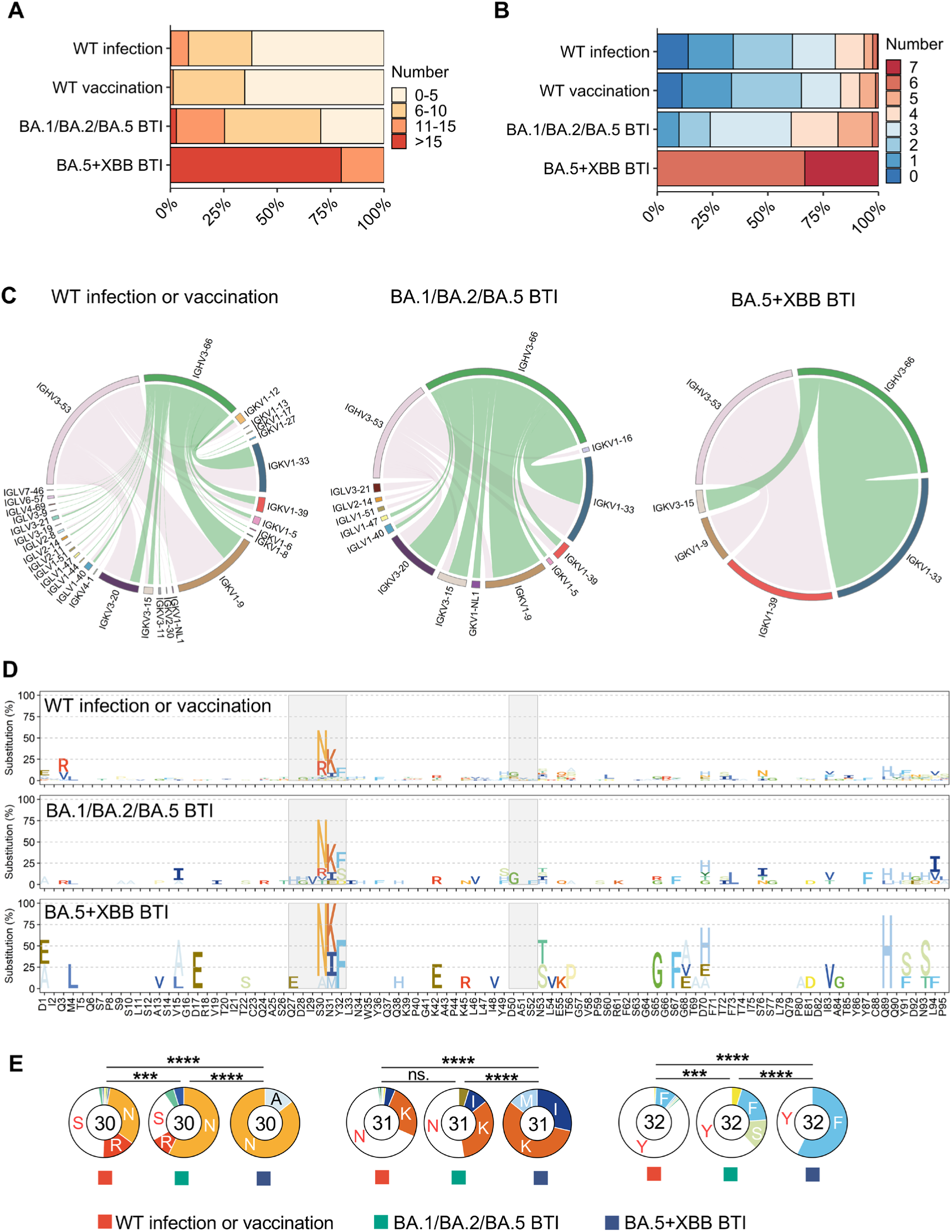
Comparison of the IGLV usage and somatic hypermutations of VH3-53/3-66 mAbs. (**A**) Comparison of the frequency distribution of total number of somatic hypermutations across VH3-53/3-66 mAbs that isolated from different infection or vaccination background. (**B**) Comparison of the frequency distribution of number of somatic hypermutations among position G26, F27, T28, S31, V50, S53, and Y58 across VH3-53/3-66 mAbs that isolated from different infection or vaccination background. (**C**) Circos plot showing the pairing light chain usage of VH3-53/3-66 mAbs that isolated from different infection or vaccination background. (**D**) Somatic mutation profiles of IGKV1-33 light chain variable region of VH3-53/3-66 mAbs that isolated from different infection or vaccination background. (**E**) Pie charts indicate the frequency of germline (white) and mutated residues at position 30-32 of IGKV1-33 light chain variable region of VH3-53/3-66 mAbs.

**Figure S4.**
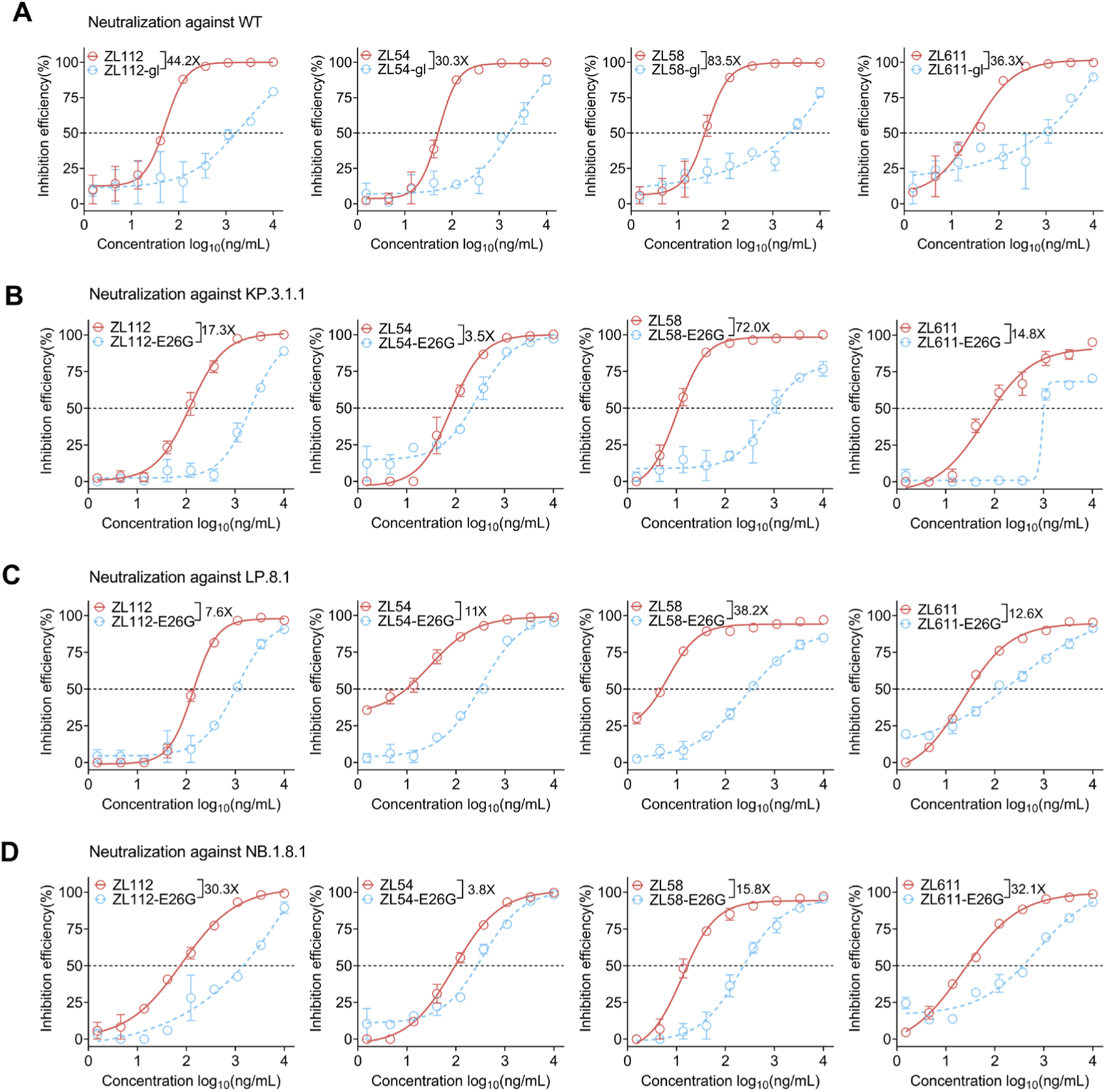
Effects of somatic hypermutation on neutralization breadth. (**A**) Neutralizing activity of prototype and germline versions of ZL112, ZL54, ZL58, and ZL611 against WT SARS-CoV-2. (**B-D**) Neutralizing activity of prototype and E26G mutant versions of ZL112, ZL54, ZL58, and ZL611 against KP.3.1.1 (**B**), LP.8.1.1 (**C**), and NB.1.8.1 (**D**).

**Figure S5.**
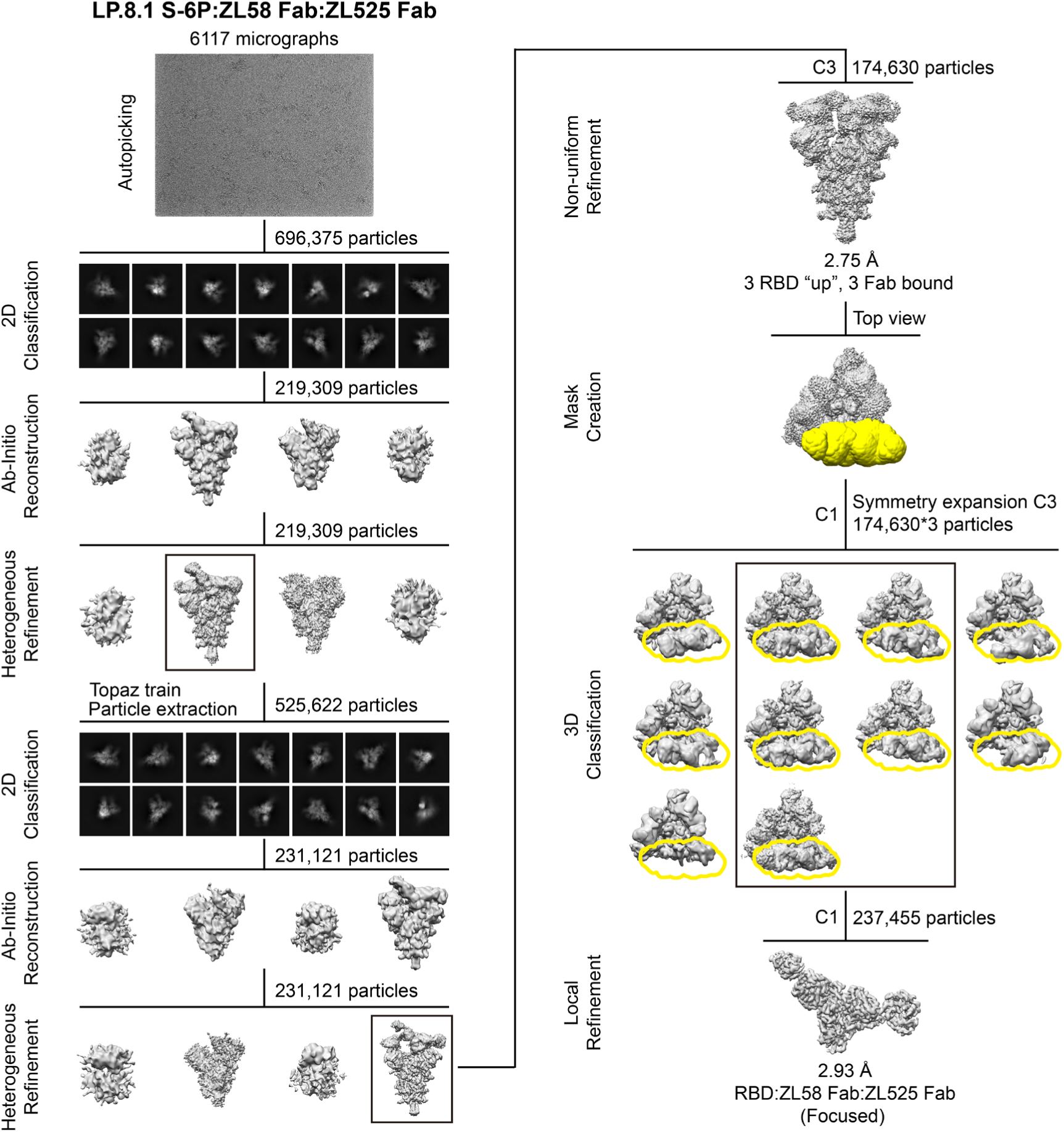
Cryo-EM data processing flow-chart for the structure of LP.8.1 S-6P:ZL58 Fab:ZL525 Fab complex. After 2D classification, ab-initio reconstruction and heterogeneous refinement were performed. High-quality particles were used to train the topaz picker, followed by particle re-extraction, 2D classification, ab-initio reconstruction, heterogeneous refinement, and non-uniform refinement. The structure of the RBD-Fab portion was further improved with focused refinement. Selected classes are highlighted with black boxes, and focused mask is shown in yellow.

**Figure S6.**
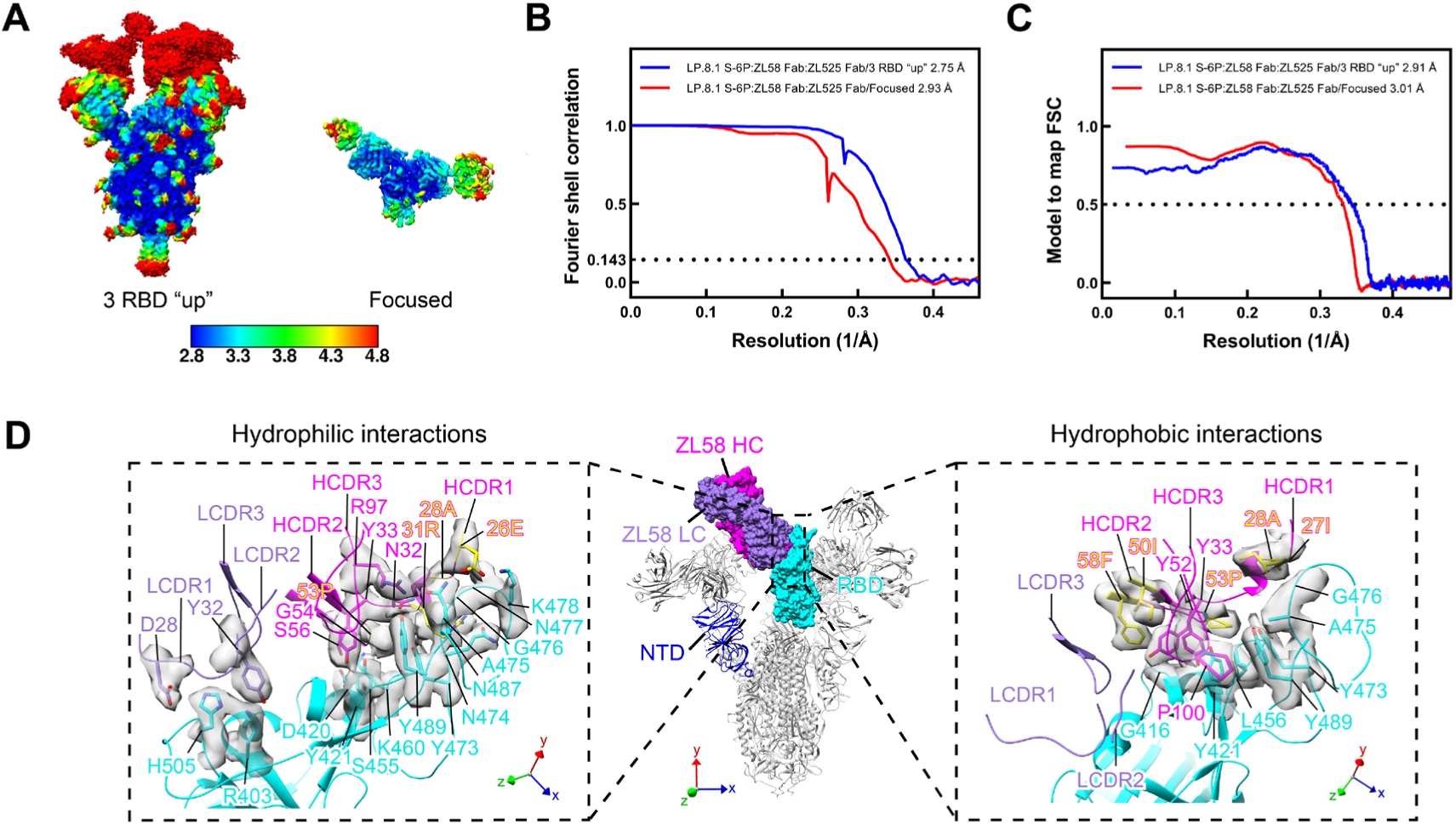
Resolution assessment of the structure of LP.8.1 S-6P:ZL58 Fab:ZL525 Fab complex. (**A**) Local resolution map for the LP.8.1 S-6P:ZL58 Fab:ZL525 Fab complex. (**B**) Global resolution assessment by Fourier shell correlation at the 0.143 criterion. (**C**) Correlations of model vs map by Fourier shell correlation at the 0.5 criterion. (**D**) Representative densities of the ZL58 epitope and ZL58 CDR loops in the focused map.

**Figure S7.**
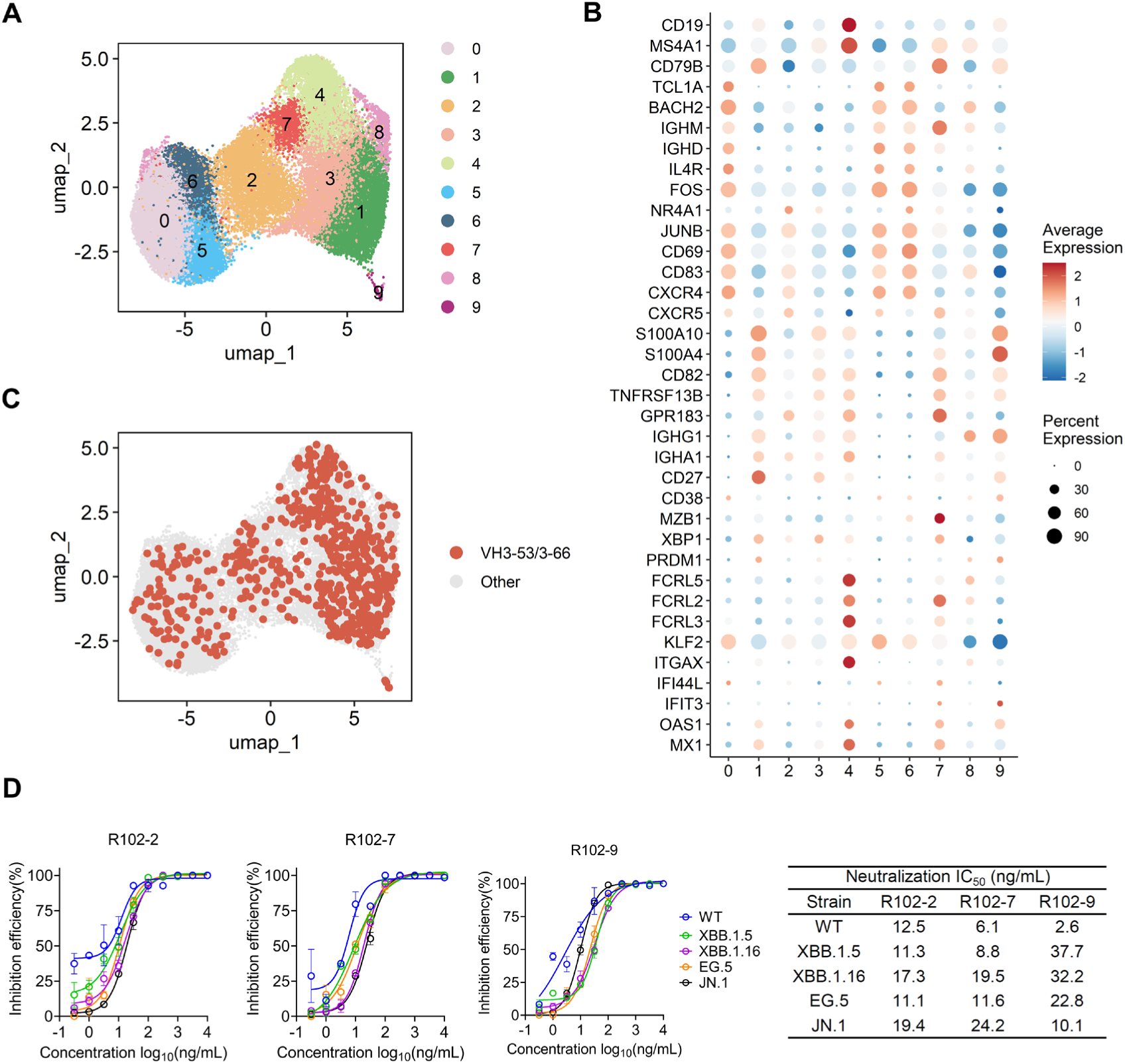
AI-based discovery of VH3-53/3-66 bnAbs from WT vaccinated donors. (**A**) Uniform manifold approximation projection (UMAP) embedding of scRNA-seq transcriptome profiles of the WT S-6P probe^+^ B cells, colored by cluster. (**B**) Dotplot showing the marker genes of the ten B cell clusters. (**C**) UMAP highlighting cells from which mAb were selected for AI virtual screening. The selected mAbs are colored with red dots and unselected mAbs with grey dots. (**D**) Authentic virus neutralizing activity of R102-2, R102-7, and R102-9 against WT, XBB.1.5, XBB.1.16, EG.5, and JN.1.

**Figure S8.**
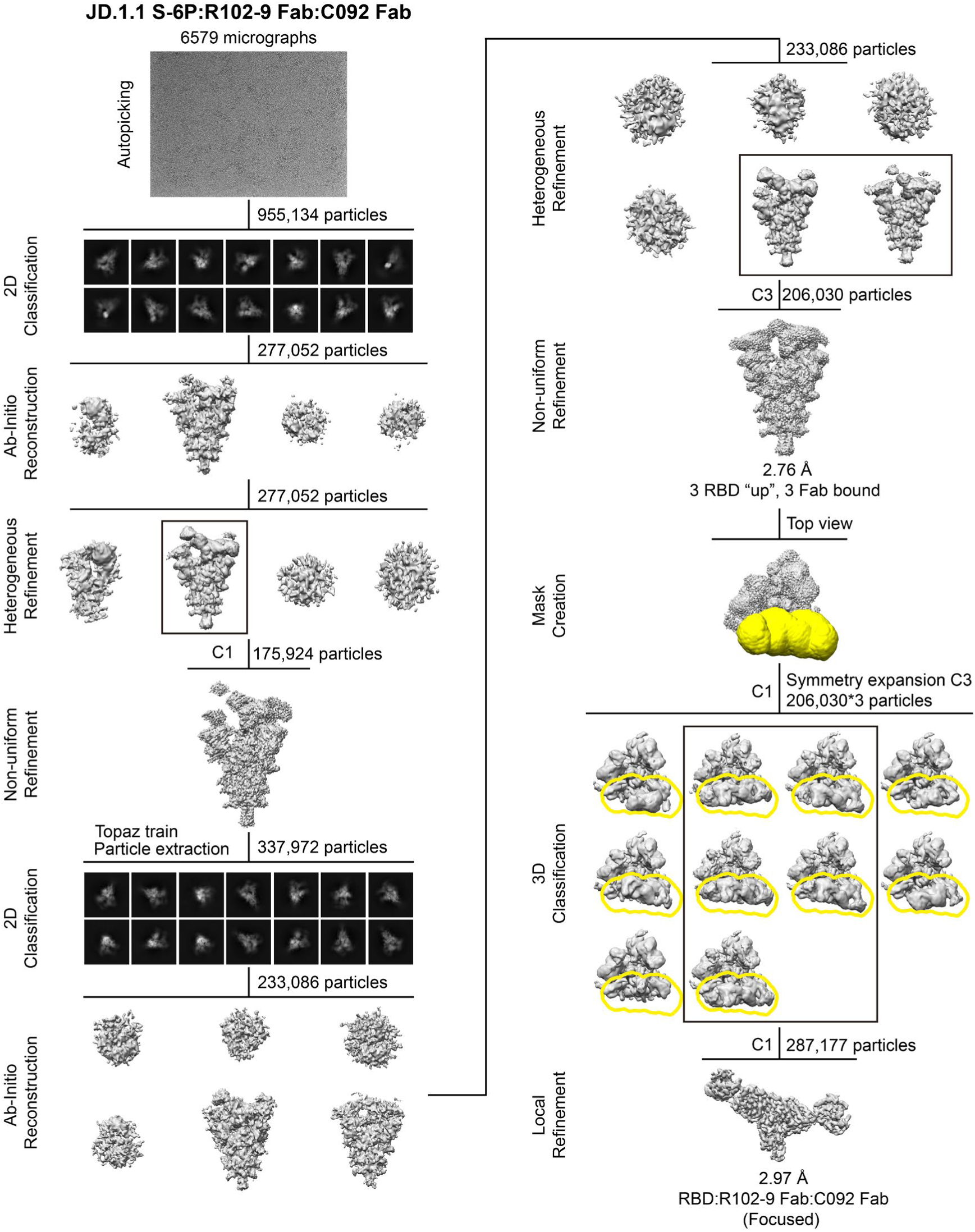
Cryo-EM data processing flow-chart for the structure of JD.1.1 S-6P:R102-9 Fab:C092 Fab complex. After 2D classification, ab-initio reconstruction, heterogeneous refinement and non-uniform refinement were performed. High-quality particles were used to train the topaz picker, followed by particle re-extraction, 2D classification, ab-initio reconstruction, heterogeneous refinement, and non-uniform refinement. The structure of the RBD-Fab portion was further improved with focused refinement. Selected classes are highlighted with black boxes, and focused mask is shown in yellow.

**Figure S9.**
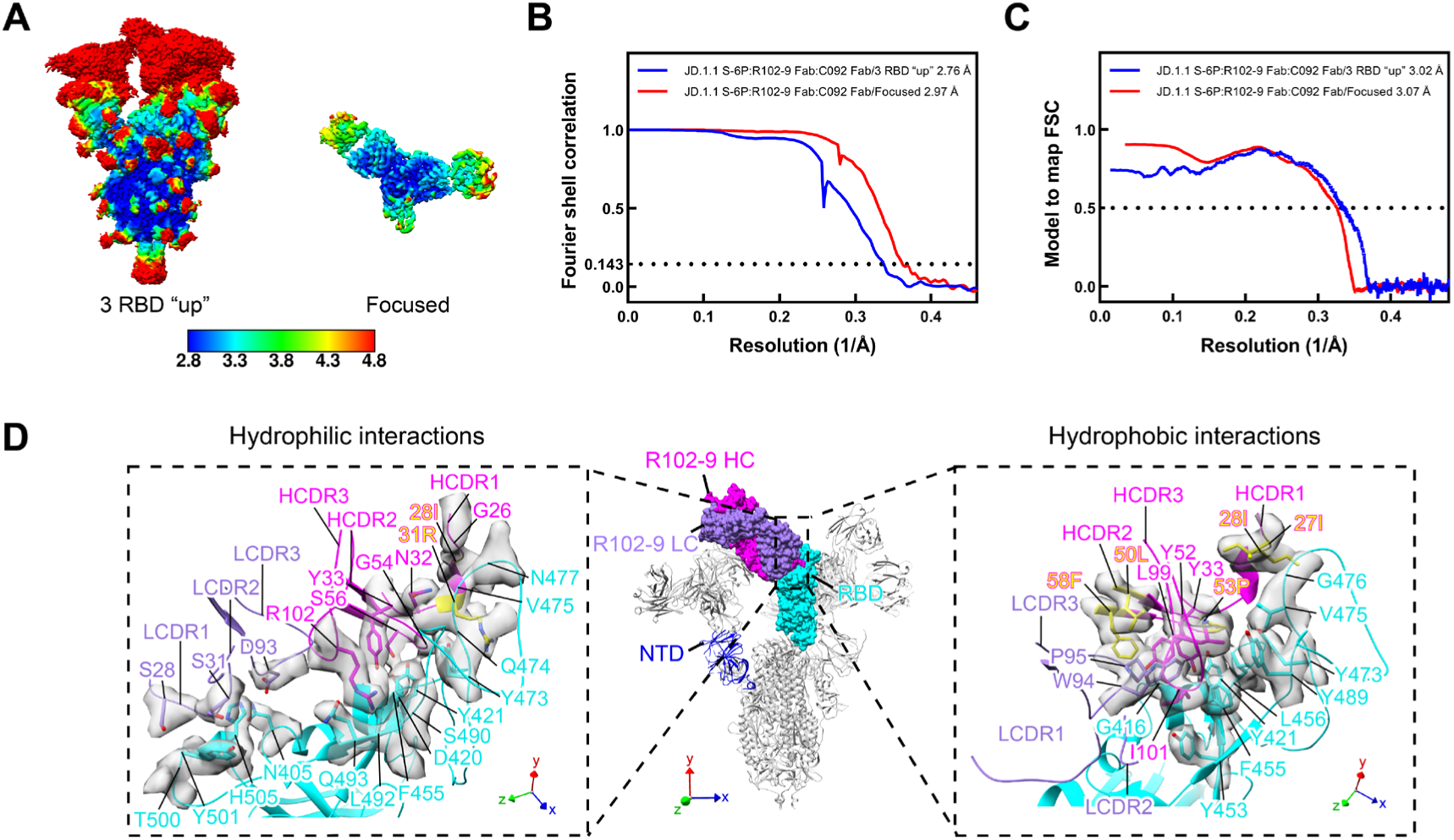
Resolution assessment of the structure of JD.1.1 S-6P:R102-9 Fab:C092 Fab complex. (**A**) Local resolution map for the JD.1.1 S-6P:R102-9 Fab:C092 Fab complex. (**B**) Global resolution assessment by Fourier shell correlation at the 0.143 criterion. (**C**) Correlations of model vs map by Fourier shell correlation at the 0.5 criterion. (**D**) Representative densities of the R102-9 epitope and R102-9 CDR loops in the focused map.

**Figure S10.**
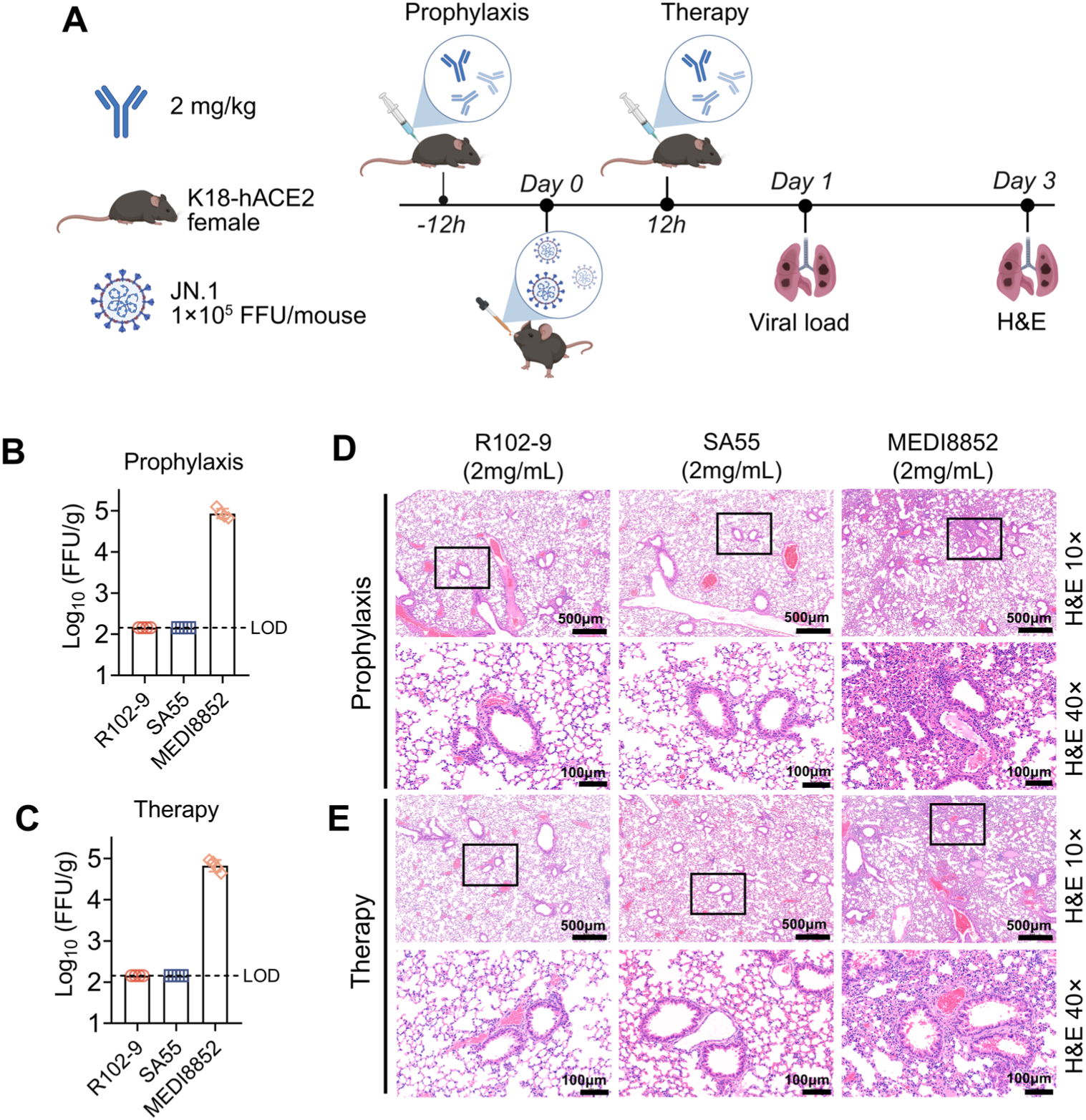
The prophylactic and therapeutic effects of R102-9 against JN.1. (**A**) Flowchart of the experimental design of testing the prophylactic and therapeutic activity of R102-9 to protect against challenge with JN.1 SARS-CoV-2 in K18-hACE2 mice. (**B-C**) Infectious virions were tested by viral fluorescent assay in lung tissue homogenates of prophylactic (**B**) and therapeutic (**C**) assay. FFUs per g of tissue extractions were compared between different groups in log_10_-transformed units. LOD, limit of detection. (**D-E**) Representative images of mouse lung tissues of prophylactic (**D**) and therapeutic (**E**) by H&E staining.

